# A computational model to study hemodynamics during atrial fibrillation

**DOI:** 10.1101/2025.06.13.659399

**Authors:** Felix Plappert, Pim J.A. Oomen, Clara E. Jones, Emmanouil Charitakis, Lars O. Karlsson, Pyotr G. Platonov, Mikael Wallman, Frida Sandberg

## Abstract

Atrial fibrillation (AF) is associated with reduced cardiac output, which is correlated with increased symptomatic burden and declined quality of life. Predicting hemodynamic effects of AF remains challenging due to the complex interplay of multiple contributing mechanisms. Computational modeling offers a valuable tool for simulating hemodynamics. However, existing models are lacking the capabilities to both replicate beat-to-beat hemodynamic variations during AF while being well suited for fitting to clinical data. In this study, we present a computational model comprising: 1) an electrical subsystem that generates uncoordinated atrial and irregular ventricular activation times characteristic of AF, and 2) a mechanical subsystem that simulates hemodynamics using a reduced order model. The model was fitted to replicate individual hemodynamic measurements from 17 patients in the SMURF study during both normal sinus rhythm (NSR) and AF. The fitted model matched a large majority (75%) of blood pressure and intracardiac pressure measurements in both NSR and AF with absolute simulation errors well below 10 mmHg. Furthermore, a large majority of left atrial and left ventricular ejection fraction measurements during NSR were matched with absolute simulation errors well below 10%. The model consistently underestimated right ventricular diastolic pressure during NSR while overestimating right ventricular systolic and mean left atrial pressures during AF. The presented approach of modeling atrial activity in AF as uncoordinated atrial contractions, rather than no atrial contraction, achieved lower overall absolute simulation errors when fitting to individual patients. This computationally efficient model provides a platform for future investigations of patient-specific hemodynamics during AF.

## 1 Introduction

Atrial fibrillation (AF) is the most prevalent supraventricular tachyarrhythmia and constitutes a significant burden for healthcare systems world wide (Hindricks et al., 2020). Atrial fibrillation impairs the heart’s pumping function and reduces the cardiac output (Morris et al., 1965; Daoud et al., 1996), which in turn is correlated with an increase in symptomatic burden and a decline in quality of life (Klavebäck et al., 2023). In the context of AF, the rate and rhythm of cardiac activity are decisive factors for the cardiac output. Atrial fibrillation is characterized by uncoordinated atrial electrical activation that results in rapid and irregular ventricular activity. In comparison to normal sinus rhythm (NSR), AF is characterized by increased heart rate, and increased RR series variability and irregularity (Carrara et al., 2015; Akande et al., 2023). These changes to ventricular rate and rhythm affect the cardiac output. The cardiac output is a function of the heart rate following an increase-plateau-decrease relationship from low to high heart rates (Kumada et al., 1967), potentially leading to a decrease in cardiac output for the high heart rates seen in AF. Moreover, the increased variability and irregularity in ventricular activity associated with AF has been shown to reduce cardiac output irrespective of heart rate (Greenfield et al., 1968; Herbert, 1973; Naito et al., 1983; Daoud et al., 1996; Clark et al., 1997). Clinical guidelines for rate control in AF only specify a target for heart rate *<*110 bpm (Van Gelder et al., 2024; Joglar et al., 2024), but it is unclear whether the impact of changes in RR series variability and irregularity relative to the impact of changes in heart rate should be accounted for in AF treatment selection. In addition to effects associated with the ventricular rhythm, the cardiac output has been shown to decrease in the absence of atrial contraction, both at a constant ventricular rate (Skinner et al., 1964) and in AF (Mitchell and Shapiro, 1969). Conversely, in response to cardioversion from AF to NSR, the cardiac output has been shown to increase, albeit with considerable variation between patients (Upshaw, 1997; Klavebäck et al., 2023). Given the complexity and multiple mechanisms involved in hemodynamic changes during AF, a means of predicting the hemodynamic effects of therapies targeting heart rate and rhythm might be valuable for AF treatment personalization.

Prediction of patient-specific hemodynamics in response to changes in heart rate and rhythm is challenging with currently available methods. A key limitation of existing computational models is their difficulty in integrating electrophysiology and hemodynamics during AF over different spatio-temporal scales. For a model to be able to replicate patient-specific hemodynamics in AF, three important criteria need to be met: 1) it includes a description of atrial and ventricular activation times characteristic of AF, 2) it is able to perform continuous hemodynamic simulations over many heartbeats to capture the beat-to-beat variation, and 3) it is efficient enough to simulate many realizations to enable patient-specific model parameter fitting. Finite element models coupled to lumped system models of the circulation system have been used to simulate cardiac hemodynamics (Kerckhoffs et al., 2007; Moyer et al., 2015; Hirschvogel et al., 2017; Augustin et al., 2021; Gerach et al., 2021) and cardiac electrophysiology in AF (Göktepe et al., 2010; Sakata et al., 2024). However, these models are computationally expensive and are typically not suited for simulations of many heart beats. Reduced order models of cardiac mechanics, such as the model by Lumens et al. (2009), offer the low computational effort necessary for replicating patient-specific hemodynamics in AF. A revision by Walmsley et al. (2015) provided a framework for spatial resolution in electrical activation patterns by dividing the cardiac walls into multiple patches. Furthermore, this model has been fitted to experimental data (Jones and Oomen, 2025). However, this family of reduced order models still lack an electrophysiological description of uncoordinated and irregular activation times characteristic of AF, which is required for studying the connection between characteristics of the atrial and ventricular rhythms and the hemodynamic performance of the heart.

The aim of the present study is to develop a simplified computational model that 1) can replicate the rapid and uncoordinated atrial activity in AF, 2) can replicate the ventricular rate and rhythm in AF characterized by heart rate, RR series variability and irregularity, 3) can be fitted to clinical data, and 4) is computationally efficient enough to allow model fitting and account for hemodynamic differences in response to rhythm by simulating a sequence of heartbeats. We extended a previously proposed hemodynamic model (Oomen et al., 2022) with an electrophysiological description that produces realistic atrial and ventricular activation times characteristic of AF and adapted the description of the atrial walls to replicate the fibrillating atria. The model was evaluated by fitting it to the SMURF dataset, described in Section 2.1, comprising clinical data collected during NSR and AF from AF patients admitted for pulmonary vein isolation (PVI). In Section 2.2, the processing of intracardiac electrogram (EGM) and electrocardiogram (ECG) signals to extract synchronized time series of atrial and ventricular activations is described. In Section 2.3, the computational model comprising an electrical subsystem (Section 2.3.1) and a mechanical subsystem (Section 2.3.2) for simulating hemodynamics in AF is described. In Section 2.4, the model fitting to replicate the patient-specific hemodynamic measurements from Section 2.1 is described. In Section 3.1, patient-specific model fits were evaluated using the entire SMURF dataset and were also presented for each patient individually. In Section 3.2, a direct comparison of patient-specific fits is presented for three distinct atrial contraction patterns that describe the atrial activity in AF: the proposed model, no atrial contraction, and regular synchronized atrial contraction.

## 2 Methods

### 2.1 Clinical data from the SMURF study

Data from a subset of the SMURF study were used in this study (Charitakis et al., 2015). Patients eligible for catheter ablation of AF with pulmonary vein isolation (PVI) were monitored using a thumb ECG device for four days before the procedure. Those who had no AF episodes during this period were included in the SMURF-randomized substudy (Charitakis et al., 2016). Twenty-nine patients were assigned to an intervention group in which AF was induced using burst pacing from the coronary sinus (CS) with a cycle length of 170-300 ms, maintained for 30 min, and re-induced immediately when necessary. Hemodynamic measurements were collected during both NSR and 30 minutes after the induction of AF. Only patients of the intervention group were considered in this study, out of which 12 individuals were excluded because of missing pressure or ejection fraction measurements. The remaining 17 patients included in the present study had a median age of 58 years (32 to 68 years) and 5 (29%) were female.

Up to 9 days before PVI, a transthoracic echocardiographic examination (TTE) was performed during NSR providing measurements of maximum left atrial volume (LAV_max_) and maximum right atrial volume (RAV_max_), and measurements of ejection fraction (EF) in the left atrium (LAEF) and the left ventricle (LVEF) (Charitakis et al., 2016) (Table 1). As part of the PVI procedure, all patients were catheterized according to clinical routine. Pressure, EGM, and ECG measurements were collected during both NSR and after 30 minutes of induced AF. Intracardiac pressures were recorded in the left atrium (LA), right atrium (RA), and right ventricle (RV) during quiet breathing for at least 15 seconds with a sampling rate of 2000 Hz and a pressure resolution of 0.5 mmHg (EP-WorkMate; St Jude Medical, Saint Paul, Minnesota, USA). From the intracardiac pressure signals, mean left atrial pressure (LAP_avg_), mean right atrial pressure (RAP_avg_), right ventricular systolic pressure (RVSP), and right ventricular diastolic pressure (RVDP) were determined. Furthermore, intracardiac EGMs were recorded in the CS using a 10-polar catheter with a sampling rate of 2000 Hz and a voltage resolution of 78 nV/LSB. Alongside intracardiac measurements, the average of three repeated non-invasive measurements of systolic blood pressure (SBP) and diastolic blood pressure (DBP) were taken during both NSR and AF (Philips Easycare) (Almroth et al., 2023). The heart rate was assessed using a 12-lead ECG. The measurements are summarized in Table 1.

**Table 1:** Volume, ejection fraction, intracardiac pressure, arterial pressure, and heart rate measurements in the study cohort in NSR and 30 min after AF induction.

| Variables | NSR | 30 min after induction of AF |
| --- | --- | --- |
| LAV<sub>max</sub> (mL) | 54 [36, 94] | – |
| RAV<sub>max</sub> (mL) | 41 [21, 64] | – |
| LAEF (%) | 53.9 [41.7, 68.3] | – |
| LVEF (%) | 64.0 [54.0, 71.0] | – |
| LAP<sub>avg</sub> (mmHg) | 15 [6, 25] | 15 [9, 19] |
| RAP<sub>avg</sub> (mmHg) | 11 [4, 17] | 11 [5, 17] |
| RVSP (mmHg) | 33 [23, 46] | 29 [17, 39] |
| RVDP (mmHg) | 9 [4, 15] | 7 [3, 14] |
| SBP (mmHg) | 127 [100, 163] | 127 [101, 175] |
| DBP (mmHg) | 79 [57, 107] | 84 [64, 115] |
| HR (bpm) | 54 [40, 81] | 95 [70, 132] |

Measurements from 17 patients are presented as median [min, max]. DBP, diastolic blood pressure; HR, mean heart rate; LAEF, left atrial ejection fraction; LAP<sub>avg</sub>, mean left atrial pressure; LAV<sub>max</sub>, maximum left atrial volume; LVEF, left ventricular ejection fraction; RAP<sub>avg</sub>, mean right atrial pressure; RAV<sub>max</sub>, maximum right atrial volume; RVDP, right ventricular diastolic pressure; RVSP, right ventricular systolic pressure; SBP, systolic blood pressure.

### 2.2 ECG and EGM signal processing

In addition to the hemodynamic measurements listed in Table 1, which were available for all 17 patients in NSR and AF, a series of atrial and ventricular activation times during AF could be extracted for 4 out of 17 patients from ECG and intracardiac EGM signals recorded during catheterization. To model patient-specific ventricular activations during AF, a series of clinical ventricular activation times ***ν***_*C*_ was extracted from ECG lead V1. The clinical ventricular activation times were estimated by the R-wave locations in the ECG. For the ECG preprocessing and R-wave detection, the free ECGdeli toolbox was used (Pilia et al., 2021). For hemodynamic simulations with a patient-specific heart rhythm (Section 2.3), a period comprising the first 200 consecutive ventricular activations in ***ν***_*C*_ of sufficient quality was selected.

Within the same time period selected for ***ν***_*C*_, atrial activation times were obtained from EGM signals. The EGM signals of the bipolar lead pair CS9-10 were used, since it was deemed to be placed closest to the entrance of the atrioventricular (AV) node. The bipolar EGM signals were preprocessed according to (Alcaine et al., 2017): 1) bandpass filtering between 40 and 250 Hz, using a second order Butterworth IIR filter; 2) signal rectification; 3) lowpass filtering with 25 Hz cut-off frequency, using a second order Butterworth IIR filter. A series of clinical atrial activation times ***α***_*C*_ was extracted from the preprocessed bipolar EGM signals using the MSPTD algorithm (Bishop and Ercole, 2018).

### 2.3 Simplified model of hemodynamics in atrial fibrillation

The simplified model of hemodynamics in AF is illustrated in Figure 1. The model is composed of an electrical subsystem (Section 2.3.1) generating atrial and ventricular activation times characteristic of AF and a mechanical subsystem (Section 2.3.2) describing the cardiovascular mechanics and hemodynamics. The electrical subsystem is proposed for the present study and provides the mechanical subsystem with atrial and ventricular activation sequences mimicking the electrical activation of the heart during AF. At the core of the electrical subsystem is an AV node network model that was previously shown to produce realistic RR series that are characteristic of AF (Karlsson et al., 2021). The cardiac electrical activity is modeled as an open loop controller, where the electrical activations drive wall contractions in the mechanical subsystem. Hence, the simulated hemodynamics influence the mechanical behavior but have no feedback effect on the electrical activations. At the core of the mechanical subsystem is a description of the heart chambers based on the TriSeg model using the MultiPatch approach (Arts et al., 2005; Lumens et al., 2009; Walmsley et al., 2015). The mechanical subsystem was previously described in Oomen et al. (2022), with the exception of the added atria divided into 20 patches to replicate the uncoordinated and irregular atrial electrical activity in AF.

**Figure 1:**
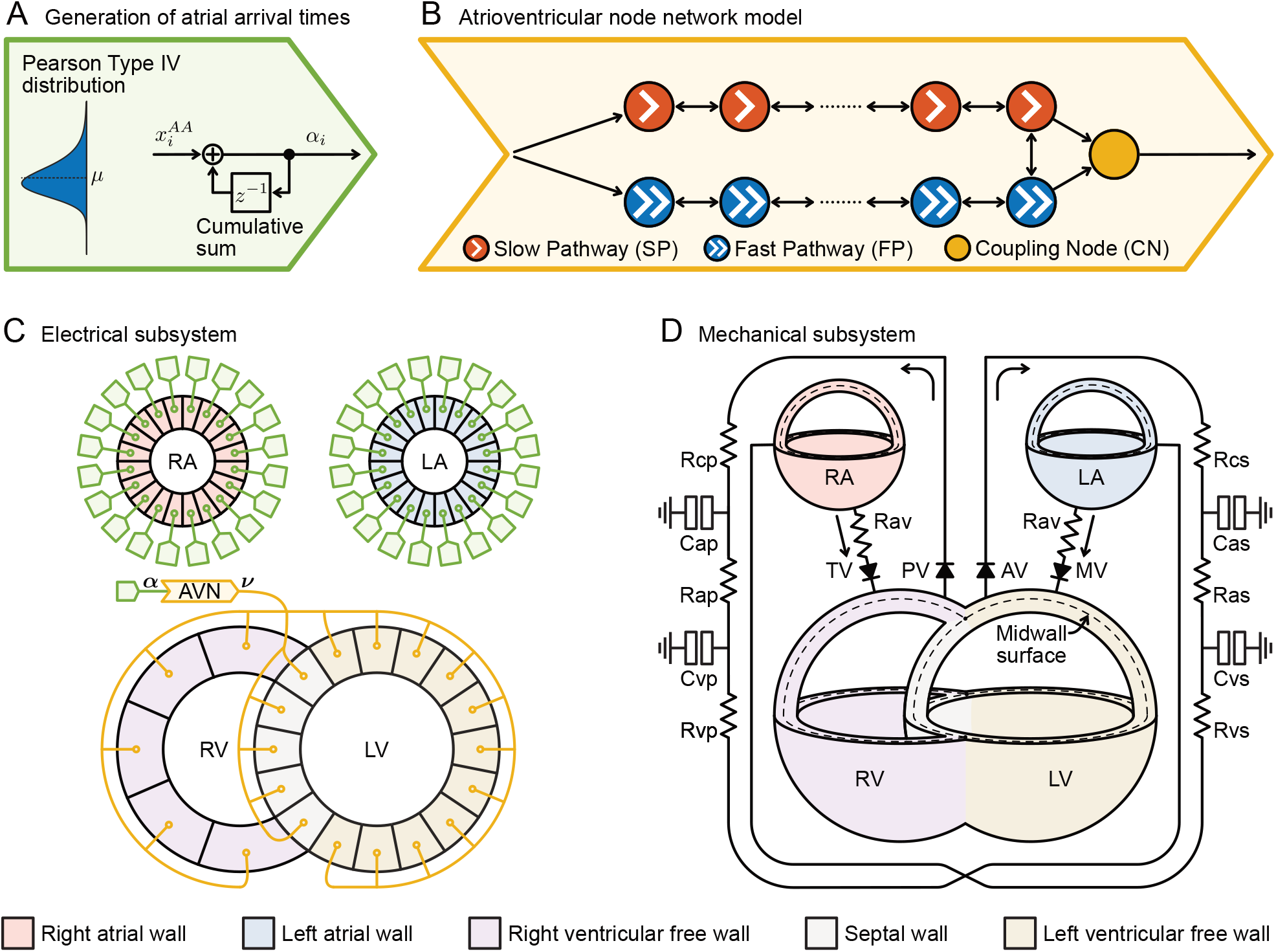
Illustration of the simplified model of hemodynamics in atrial fibrillation. The model comprises a subsystem describing cardiac electrical activation times in atrial fibrillation (A-C) and a subsystem describing the cardiovascular mechanics and hemodynamics (D). **A)** A series of atrial activation times ***α*** was computed as the cumulative sum of independent samples from a Pearson Type IV distribution. **B)** The AV node was modeled as a network of nodes and produced a series of ventricular activation times ***ν*** = ***ν***_*S*_. **C)** Each atrial wall patch and the AV node model were activated by a unique series of atrial activation times ***α***. All ventricular wall patches were activated by the same series ***ν*. D)** The heart chambers were described based on the TriSeg model using the MultiPatch approach. The heart valves and circulation system were described equivalent to an electrical circuit.

#### 2.3.1 Electrical subsystem generating cardiac electrical activation times in atrial fibrillation

To reproduce the uncoordinated and irregular atrial activity characteristic of AF, a unique series of atrial activation times was generated for each atrial wall patch. A wall patch describes a collection of cells in the cardiac wall that share the same mechanical properties and electrical activation time (see Section 2.3.2 for more detail). For the generation of atrial activation times (Figure 1A), a series of atrial-to-atrial activation intervals 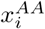 was independently drawn from a Pearson Type IV distribution with mean *µ*, standard deviation *σ*, skewness *γ*, and kurtosis *κ* (Climent et al., 2011). *A series* ***α*** of atrial activation times *α*_*i*_ was then obtained using the cumulative sum of the series of AA intervals, defined in Eq. 1 as

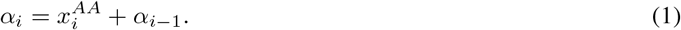

In all 17 patients, ***α*** = ***α***_*S*_ was produced with *µ* randomly drawn from 𝒰 [100, 250]ms, *σ* randomly drawn from 𝒰 [15, 30]ms, and *γ* and *κ* fixed at 1 and 6, respectively (Plappert et al., 2022). Additionally, for the 4 patients where clinical EGM signals were available, 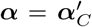 was produced with *µ, σ, γ*, and *κ* extracted from ***α***_*C*_ described in Section 2.2.

The AV node was modeled by a network of 21 nodes (Figure 1B) (Wallman and Sandberg, 2018; Karlsson et al., 2021). The slow pathway (SP) and fast pathway (FP) are described by two chains of 10 nodes each, connected only at their last nodes. Impulses enter the AV node model simultaneously at the first node of each pathway and can travel bidirectionally between connected nodes. The last nodes of the two pathways have a unidirectional connection to an additional coupling node (CN), through which impulses exit the model.

The AV node model processes a series of atrial impulses chronologically and node by node, using a priority queue of nodes, sorted by impulse arrival time (Wallman and Sandberg, 2018). Input to the network AV node model was a unique series of atrial activation times ***α*** generated using the same Pearson Type IV distribution as the activation times for the atrial wall patches (Figure 1C). Impulses arriving at a node can be either conducted to all adjacent nodes with a delay or blocked due to refractoriness. An impulse is conducted if Δ*t*_*k*_ *≥* 0 and blocked if Δ*t*_*k*_ *<* 0, where Δ*t*_*k*_ is the interval between the *k*:th conducted impulse arrival time *t*_*k*_ and the end of the (*k* − 1):th refractory period *R*^*P*^(Δ*t*_*k*−1_), as defined in Eq. 2

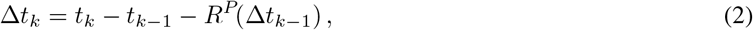

where *P ∈* {*SP, FP, CN*} denotes the pathway. If conducted, the arrival of the *k*:th impulse to all adjacent nodes is delayed by the conduction delay *D*^*P*^(Δ*t*_*k*_). Each node is characterized by an individual refractory period *R*^*P*^(Δ*t*_*k*_) and conduction delay *D*^*P*^(Δ*t*_*k*_) defined in Eqs. 3, 4 as

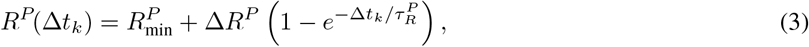

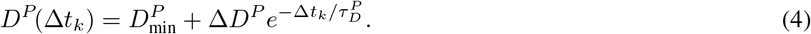

All nodes of a pathway *P* were characterized by the same six fixed parameters 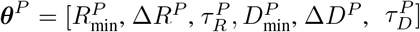. The parameters of the SP and FP were drawn from bounded uniform distributions, and the model parameters of the CN were kept fixed according to Table 2, as previously proposed in (Plappert et al., 2022). Some of the drawn parameter sets will be excluded based on the selection criteria described in Section 2.4.1, resulting in non-uniform parameter distributions for the SP and FP. The output of the network AV node model is a series of ventricular activation times ***ν***_*S*_ containing all *t*_*k*_ of the CN.

**Table 2:** AV node model parameters used for simulated data.

| Parameters | P≡SP (ms) | P≡FP (ms) | P≡CN (ms) |
| --- | --- | --- | --- |
| $R_{\min}^P$ | $\mathcal{U}[250, 600]$ | $\mathcal{U}[250, 600]$ | 250 |
| $\Delta R^P$ | $\mathcal{U}[0, 600]$ | $\mathcal{U}[0, 600]$ | 0 |
| $\tau_R^P$ | $\mathcal{U}[50, 300]$ | $\mathcal{U}[50, 300]$ | 1 |
| $D_{\min}^P$ | $\mathcal{U}[0, 30]$ | $\mathcal{U}[0, 30]$ | 0 |
| $\Delta D^P$ | $\mathcal{U}[0, 75]$ | $\mathcal{U}[0, 75]$ | 0 |
| $\tau_D^P$ | $\mathcal{U}[50, 300]$ | $\mathcal{U}[50, 300]$ | 1 |

To reproduce the coordinated but irregular ventricular activity characteristic of AF, the same series of ventricular activation times ***ν*** was used for all ventricular wall patches in a simulation (Figure 1C). In all 17 patients, ***ν*** = ***ν***_*S*_ with 200 ventricular activation times was produced by the AV node network model. Additionally, for the 4 patients where clinical EGM signals were available, ***ν*** = ***ν***_*C*_ was extracted from the ECG signal as described in Section 2.2.

#### 2.3.2 Mechanical subsystem describing the cardiovascular mechanics and hemodynamics

The hemodynamics in NSR and AF were simulated using a previously published mechanical model (Oomen et al., 2022) (Figure 1D). The description of the heart chambers was based on the Three-Wall Segment (TriSeg) method (Lumens et al., 2009) with atrial and ventricular walls divided into multiple segments using the MultiPatch module (Walmsley et al., 2015). The heart valves and circulation system were described as equivalent to an electrical circuit, with heart valves modeled as diodes, and the systemic and pulmonary circulation modeled as 5-element RCRCR Windkessel model (Figure 1D) (Santamore and Burkhoff, 1991). Using the mechanical subsystem, the pressure and volume dynamics within the heart chambers and circulation system were simulated as a closed loop system with a time step of 0.5 ms. The change in blood volume between heart chambers and vasculature was computed using a 4^th^ order Runge-Kutta solver. The total blood volume was functionally divided into stressed blood volume (SBV) and unstressed blood volume, with the latter being the volume that fits into the cardiovascular system before pressure rises above 0 mmHg. According to the TriSeg method (Lumens et al., 2009), the ventricles were modelled as thick-walled spherical segments comprising the left ventricular free wall (LFW), right ventricular free wall (RFW), and septal wall (SW) interconnected at a circular wall junction. The atria were modeled as separate thick-walled spheres. No interaction between the atria and ventricles was assumed, due to the presence of the dense connective tissue separating these chambers. The description of heart valves as pressure-sensitive diodes allowed unidirectional blood flow from the atria to the ventricles and from the ventricles to the arteries. Heart valve regurgitation was not accounted for in the present study.

The spherical wall segments were divided into multiple patches using the MultiPatch approach (Walmsley et al., 2015). The LFW and SW consisted of 11 and 5 wall patches, respectively, corresponding to the 16 AHA segment norm for the left ventricle (Schiller et al., 1989). The RFW consisted of 5 wall patches as previously modelled in Oomen et al. (2022). For the present study, the left atrial wall (LAW) and right atrial wall (RAW) each consisted of 20 wall patches to allow modelling of uncoordinated and irregular atrial activity in AF. Each wall patch was activated independent from other wall patches; this activation was initiated by the electrical subsystem, described in Section 2.3.1. The cardiac mechanics were calculated at the midwall surface area of each patch, dividing each thick-walled spherical patch into two equal volumes (Figure 1D). Myofiber strain was derived from the midwall surface area in relation to a reference area *A*_m,ref_, as described in Walmsley et al. (2015). Total Cauchy myofiber stress was calculated as the sum of an active and passive stress component. The active stress was calculated using a Hill-type sarcomere active contraction model and depended on how much the wall patch was stretched and on mechanical activation, and contained a linear scaling factor for active myofiber stress *S*_act_ and an exponential scaling factor for the duration of contraction *t*_ad_ (Lumens et al., 2009). The passive stress was calculated using an exponential constitutive model and depended on the myofiber strain (Oomen et al., 2022). The midwall tension of an entire cardiac wall segment was calculated from the myofiber stress that was allowed to vary between wall patches as described in Walmsley et al. (2015). To satisfy the principle of conservation of energy, a mechanical equilibrium of tensile forces at the wall junction between LFW, RFW and SW is maintained. Each time step, an iterative Newton scheme was employed to adjust the curvature and distance between the wall junctions of LFW, SW, and RFW until the midwall tension at the circular wall junction was below 1 *µ*N. When simulating the hemodynamics, the pressures in all compartments were updated based on the current midwall tensions and curvatures. The updated compartment pressures determine the midwall surface area, used for computing myofiber strain, stress, and midwall tension of the next iteration. The model parameters of the mechanical subsystem are listed in Tables Appendix, A2, and A3.

### 2.4 Patient-specific model fitting

For the model fitting, hemodynamic simulations were performed with patient-specific electrical activations ***α*** and ***ν*** as described in Section 2.4.1. The initialization of the hemodynamic simulations and computation of hemodynamic indices was described in Section 2.4.2. For the model fitting, a parameter space was first iteratively reduced with bayesian history matching (BHM) utilizing Gaussian process emulators (GPEs) until all remaining solutions fell within a plausible range around the target data. Subsequently, simulations were conducted for a random selection of 1 024 parameter sets drawn from the final parameter space, from which a final model parameter set was selected based on a cost function *J*, as described in Section 2.4.3. Finally, the evaluation of the patient-specific model fit is detailed in Section 2.4.4.

#### 2.4.1 Patient-specific electrical model parameter fitting

The first step in simulating the hemodynamics was to produce a patient-specific series of atrial and ventricular activation times ***α*** and ***ν*** for every wall patch. In NSR, ventricular activations in ***ν***_NSR_ were assumed to occur at a fixed rate corresponding to the patient-specific heart rate. Further, the AV nodal conduction delay was assumed to be fixed at 120 ms resulting in ***α***_NSR_ = ***ν***_NSR_ − 120 ms. Regarding ***α***, in NSR it was assumed that the regular atrial contraction can be approximated by the whole atria contracting at once. This assumption is equivalent to reducing the number of atrial wall patches in the mechanical model from 20 to 1.

In AF, the model was fitted to either ECG-derived activation times in ***ν***_*C*_ and EGM-derived activation times in 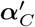 or synthetic activation times in ***ν***_*S*_ and ***α***_*S*_, resulting in two model variants. In 4 out of 17 patients, the first model variant was fitted using patient-specific ventricular activation times, ***ν***_AF_ = ***ν***_*C*_, extracted from available ECG signals as described in Section 2.2. Further, unique series 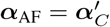 were generated for each atrial wall patch from a series of clinical activation times from a single atrial location ***α***_*C*_, as described in Section 2.3.1. The second model variant was fitted to all 17 patients using a patient-specific series of ventricular activation times, ***ν***_AF_ = ***ν***_*S*_, based on the heart rate and generated using the electrical subsystem described in Section 2.3.1. For the generation of ***ν***_*S*_, it was assumed that the interval between two adjacent ventricular activations in ***ν***_*S*_ corresponded to an RR interval. Following that assumption, an RR series was extracted from ***ν***_*S*_ and the RR mean 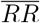, RR series variability *RR*_*V*_, and RR series irregularity *RR*_*I*_ were computed as described in Plappert et al. (2022) with the exception that *RR*_*V*_ was obtained in the present study as the root mean square of successive RR interval differences (RR rmssd) normalized with respect to 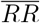. The model parameters of the electrical subsystem were randomly drawn until the following four conditions were met: 1) the SP had a lower refractory period *R*^*SP*^(Δ*t*_*k*_) *< R*^*F P*^(Δ*t*_*k*_) and higher conduction delay *D*^*SP*^(Δ*t*_*k*_) *> D*^*F P*^(Δ*t*_*k*_) than the FP for all Δ*t*_*k*_ (Plappert et al., 2022), *2)* 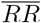 was within 1.25 bpm of the heart rate, 3) *RR*_*V*_ *∈* [0.32, 0.33] (McManus et al., 2013; Corino et al., 2015), 4) *RR*_*I*_ *∈* [1.69, 1.71] (Corino et al., 2015). In addition to generating ***ν***_*S*_, a unique series ***α***_AF_ = ***α***_*S*_ was produced for each atrial wall patch as described in Section 2.3.1.

#### 2.4.2 Patient-specific hemodynamic simulations

The second step in simulating the hemodynamics was to perform the simulations with the mechanical subsystem. The patient-specific model parameters for the mechanical subsystem were determined using the parameter fitting described in Section 2.4.3. For both NSR and AF, the simulated pressure and volume trends were segmented by cardiac cycles. A cardiac cycle was defined separately for the right and left heart by a sequence of valve events. This sequence began with the closure of the AV valve, continued with the opening of the semilunar valve, then the closure of the semilunar valve, the opening of the AV valve, and ended with the next closure of the AV valve. From each cardiac cycle, arterial blood pressure (SBP, DBP), atrial and ventricular blood pressures (LAP_avg_, RAP_avg_, RVSP, RVDP), atrial and ventricular volumes (LAV_max_, RAV_max_, left ventricular diastolic volume (LVDV), right ventricular diastolic volume (RVDV)), ejection fractions (LAEF, RAEF, LVEF, RVEF), and cardiac output (CO) were extracted. The parameter fitting utilized the mean values of these hemodynamic indices across all cycles. For certain model parameter sets and short RR intervals, the AV valve may not open and close between consecutive ventricular contractions, resulting in a saddle point in the volume trend where the ventricle does not fill between two ventricular contractions. In such cases, these consecutive ventricular contractions are treated as a single cardiac cycle for the computation of the hemodynamic indices.

Each simulation in NSR and AF was initialized with ventricular activations occuring at a fixed rate corresponding to the patient-specific heart rate and atrial activations 120 ms preceding the ventricular activations until the volumes at the start and end of the cardiac cycle differed less than 1% of the total *SBV*. Once steady state was reached, simulations in NSR began using ***α***_NSR_ and ***ν***_NSR_. With regular ventricular activation in ***ν***_NSR_, beat-to-beat hemodynamic variation is negligible; thus, the hemodynamic indices were based on one cardiac cycle. For the AF simulations, an additional AF launch phase was performed before simulating the hemodynamics to avoid transient effects. During the AF launch phase, the ventricles activate 10 times, while the atrial wall patches activate as many times as needed to match the total duration of the ventricular activations. The atrial and ventricular activation times for the AF launch phase were generated using the same model parameters that produced ***α***_AF_ and ***ν***_AF_. Then, the hemodynamics in AF were simulated using a unique series ***α***_AF_ for every atrial wall patch and a series ***ν***_AF_ with 200 ventricular activations for all ventricular wall patches as described in Section 2.4.1.

#### 2.4.3 Patient-specific mechanical model parameter fitting

Previously, BHM utilizing GPEs has been used for the calibration of the mechanical model in Jones and Oomen (2025), where a more detailed description of BHM and GPE can be found. In the present study, BHM utilizing GPEs was used to fit the combined electrical and mechanical model to match the clinical data described in Section 2.1. Parameter sets ***p***, each containing sixteen mechanical model parameters, were fitted to mimic patient-specific hemodynamics. An overview of these model parameters is given in Table 3. The size of the heart chambers was fitted by adjusting the midwall reference area *A*_m,ref,*W*_ with *W ∈* {RA, LA, RFW, SW, LFW}. Furthermore, the atrial and ventricular wall mechanics were fitted by adjusting the active stress coefficient (*S*_act,a_, *S*_act,v_) and contraction duration (*t*_ad,a_, *t*_ad,v_). The circulation system was fitted by adjusting the stressed blood volume *SBV*, the systemic and pulmonary arterial resistance *R*_as_ and *R*_ap_, and the systemic and pulmonary capacitances using a scaling factor *k*_Cs_ and *k*_Cp_ (Figure 1D). The cardiac contraction duration and the resistances and capacitances in the circulation system were allowed to change between NSR and AF to account for autonomic nervous system (ANS) induced changes during AF; these changes were accounted for by the scaling factors *k*_1_ and *k*_2_, respectively. The remaining parameters were identical between NSR and AF for each patient. The scaling factors *k*_Cs_, *k*_Cp_, *k*_1_ and *k*_2_ affect the cardiac contraction duration and the resistances and capacitances in the circulation system according to Table 3.

**Table 3:** Model parameters of mechanical subsystem fitted using Bayesian history matching.

|  | Parameter | Range | Unit | Notes |
| --- | --- | --- | --- | --- |
| Cardiac geometry | $A_{m,ref,RA}$ | [5 500, 12 500] | mm<sup>2</sup> | |
| | $A_{m,ref,LA}$ | [4 500, 8 500] | mm<sup>2</sup> | |
| | $A_{m,ref,RFW}$ | [11 000, 22 000] | mm<sup>2</sup> | |
| | $A_{m,ref,SW}$ | [3 500, 9 000] | mm<sup>2</sup> | |
| | $A_{m,ref,LFW}$ | [4 000, 11 000] | mm<sup>2</sup> | |
| Cardiac mechanics | $S_{act,v}$ | [0.1, 0.6] | MPa | |
| | $S_{act,a}$ | [0.1, 0.6] | MPa | |
| | $t_{ad,v}$ | $[75, 500] \cdot k_1^{-1}$ | ms | |
| | $t_{ad,a}$ | $[75, 500] \cdot k_1^{-1}$ | ms | |
| | $k_1$ | 1 (NSR) | – | $k_1 = 1$ during normal sinus rhythm |
| | | [0.6, 1.4] (AF) | – | $k_1 \in [0.6, 1.4]$ during atrial fibrillation |
| Circulation system | $SBV$ | [100, 2 500] | mL | |
| | $R_{as}$ | $[0.5, 2.0] \cdot k_2$ | mmHg·s·mL<sup>-1</sup> | |
| | $R_{ap}$ | $[0.02, 0.6] \cdot k_2$ | mmHg·s·mL<sup>-1</sup> | |
| | $k_{Cs}$ | [0.8, 1.2] | – | $C_{as} = 1.32 \cdot k_{Cs} \cdot k_2^{-1}$, $C_{vs} = 70 \cdot k_{Cs} \cdot k_2^{-1}$ |
| | $k_{Cp}$ | [0.8, 1.2] | – | $C_{ap} = 13 \cdot k_{Cp} \cdot k_2^{-1}$, $C_{vp} = 8 \cdot k_{Cp} \cdot k_2^{-1}$ |
| | $k_2$ | 1 (NSR) | – | $k_2 = 1$ during normal sinus rhythm |
| | | [0.6, 1.4] (AF) | – | $k_2 \in [0.6, 1.4]$ during atrial fibrillation |

For the BHM, the entire parameter space was initially sampled with 2^22^ parameter sets using a Sobol sequence (Sobol’, 1967; Joe and Kuo, 2008). To exclude non-physiological parameter sets from the parameter space, 1) *k*_1_ and *k*_2_ both had to be *≥*1 or *≤*1, and 2) the atrial contraction duration had to be lower than or equal to the ventricular contraction duration *t*_ad,a_ *≤ t*_ad,v_ (Nollet et al., 2020), resulting in *≈* 2^20^ parameter sets. To reduce computational cost, the entire parameter space was evaluated with GPEs as a surrogate model that was trained in each iteration of the BHM method on 128 simulations of our proposed model during NSR and AF. The simulations in NSR and AF were performed as described in Section 2.4.1.

Bayesian history matching is a statistical approach that can iteratively reduce a parameter space defined within conservative ranges until all remaining solutions fall within a plausible range around the target data. For every point ***p*** in the parameter space (i.e., parameter set), a maximum implausibility score *I*(***p***) was computed as defined in Eq. 5

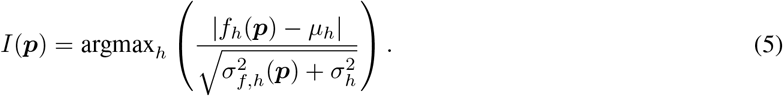

The implausibility score considered the 18 hemodynamic indices *h* listed in Table 4. Here, *f*_*h*_(***p***) is the emulated value, *µ*_*h*_ is the target mean, and 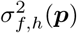 and 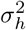 are the variance of the emulations and the target variance, respectively. An emulation *f*_*h*_(***p***) was produced for every hemodynamic index *h* and every parameter set ***p***. Furthermore, for each hemodynamic index *h*, the variance 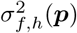 of the emulations *f*_*h*_(***p***) was computed over all ***p***. A parameter set ***p*** remained in the parameter space at the end of a BHM iteration if *I*(***p***) was below a threshold *I*_threshold_ (Vernon et al., 2010; Craig et al., 1997). The BHM procedure was executed a minimum of 7 and a maximum of 20 iterations until convergence was achieved. Between iterations 1 and 7, the threshold *I*_threshold_ was progressively reduced from 5 to 2 in decrements of 0.5. The BHM converged when *≥*90% of the points emulated in an iteration had an implausibility score *I*(***p***) *≤* 2. A threshold *I*_threshold_ = 2 guarantees that the interval [*µ*_*h*_ − 2*σ*_*h*_, *µ*_*h*_ + 2*σ*_*h*_] contains at least 95% of the hemodynamic indices *h* that result from the remaining parameter sets in the parameter space, corresponding to a 95% confidence interval for the target data, where *µ*_*h*_ and *σ*_*h*_ are listed in Table 4. For pressures and ejection fractions, *µ*_*h*_ were set to the patient-specific hemodynamic measurements summarized in Table 1, whereas *σ*_*h*_ were equal for all patients as summarized in Table 4 and calculated as described in the Appendix. Because volume measurements collected with TTE are underestimated (Whitlock et al., 2010; Kühl et al., 2012; Kitano et al., 2019), *µ*_*h*_ and *σ*_*h*_ for volumes were set to sex-specific ranges from cardiac computed tomography angiography measurements in literature (Fuchs et al., 2016), rather than to the patient-specific measurements in Table 1. To maintain a parameter space with at least 100 000 points, additional parameter sets were generated at the end of an iteration using the cloud method (Coveney and Clayton, 2018).

**Table 4:** Hemodynamic indices *h* accounted for in the BHM based fitting with sex-specific values for volumes, where M is male and F is female. ‘patient-specific’ refers to the hemodynamic index *h* of the individual patient.

| Variable $i$ | $\mu_h$ (in NSR) | $\mu_h$ (in AF) | $\sigma_h$ (in NSR, AF) |
| --- | --- | --- | --- |
| SBP (mmHg) | patient-specific | patient-specific | 20.47 |
| DBP (mmHg) | patient-specific | patient-specific | 20.47 |
| RVSP (mmHg) | patient-specific | patient-specific | 8.75 |
| RVDP (mmHg) | patient-specific | patient-specific | 8.75 |
| LAP<sub>avg</sub> (mmHg) | patient-specific | patient-specific | 6.92 |
| RAP<sub>avg</sub> (mmHg) | patient-specific | patient-specific | 6.92 |
| LAEF (%) | patient-specific | – | 5 |
| LVEF (%) | patient-specific | – | 5 |
| LAV<sub>max</sub> (mL) | M: 81, F: 67 | – | M: 18, F: 14 |
| RAV<sub>max</sub> (mL) | M: 106, F: 81 | – | M: 26, F: 18 |
| LVDV (mL) | M: 133, F: 105 | – | M: 28, F: 19 |
| RVDV (mL) | M: 173, F: 130 | – | M: 36, F: 24 |

DBP, diastolic blood pressure; LAEF, left atrial ejection fraction; LAP<sub>avg</sub>, mean left atrial pressure; LAV<sub>max</sub>, maximum left atrial volume; LVEF, left ventricular ejection fraction; RAP<sub>avg</sub>, mean right atrial pressure; RAV<sub>max</sub>, maximum right atrial volume; RVDP, right ventricular diastolic pressure; RVDV, right ventricular diastolic volume; RVSP, right ventricular systolic pressure; RVDV, right ventricular diastolic volume; SBP, systolic blood pressure.

For all patients, simulations were conducted using a random selection of 1 024 parameter sets from the final parameter space once convergence was achieved in the BHM method. As a last step in the parameter fitting procedure, a final model parameter set was selected from these parameter sets by minimizing a cost function *J* (Eq. 6) based on patient-specific fits of arterial and intracardiac pressures, ejection fractions and atrial volumes.

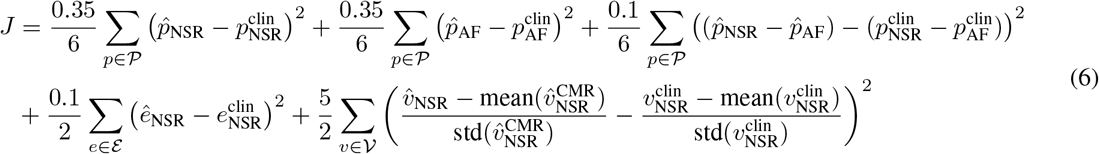

The first two terms considered the squared difference between simulated 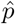 and clinical *p*^clin^ pressures 𝒫 *∈* {SBP, DBP, RVSP, RVDP, LAP_avg_, RAP_avg_} during NSR and AF. The third term considered the squared pressure changes between NSR and AF. The fourth term accounted for the squared differences between simulated *ê* and clinical *e*^clin^ left atrial and left ventricular ejection fractions ℰ *∈* {LAEF, LVEF} in NSR. The fifth term considered the squared relative size of atrial volumes 𝒱 *∈* {LAV_max_, RAV_max_} in NSR using simulated 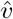, clinical TTE *v*^clin^, and reference cardiac magnetic resonance (CMR) *v*^CMR^ volumes. The 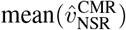 and 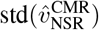 denote the average and standard deviation of atrial CMR volume measurements in a study cohort of 569 patients (Fuchs et al., 2016). The approach of relating individual volume measurements and simulations to the cohort distributions was chosen to account for systematic differences between TTE- and CMR-based volume measurements while retaining information from the TTE-based clinical measurements. To illustrate the predictive uncertainty of the BHM, a posterior distribution was produced for the hemodynamic indices using each of the 1 024 parameter sets that resulted in *I*(***p***) *≤* 2.

#### 2.4.4 Evaluation of patient-specific model fit

As described in Section 2.4.1, the first model variant using 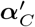 and ***ν***_*C*_ was fitted to 4 patients with available ECG and EGM signals, whereas the second model variant using ***α***_*S*_ and ***ν***_*S*_ was fitted to all 17 patients. For the evaluation of the model fit, one model per patient was accounted for in the statistics. In the case where 4 patients had two model variants, the variant using 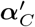 and ***ν***_*C*_ was always chosen because the fitting incorporated more clinical measurements. Each model was evaluated based on the hemodynamic indices that were computed using the final model parameter set from the BHM, selected with the cost function *J* defined in Eq. 6. For each hemodynamic index in the set 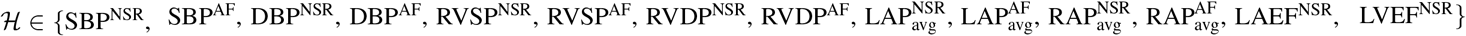, the model fit was evaluated using the simulation error *ϵ*_*ℋ*_ and normalized simulation error *ϵ*_*ℋ*,norm_ defined in Eqs. 7, 8 as

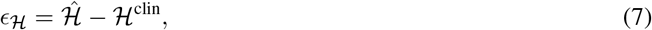

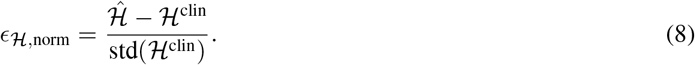

The two errors considered simulated indices 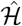 and clinical measurements ℋ^clin^. The normalized simulation error *ϵ*_ℋ,norm_ was scaled by the standard deviation of the clinical measurement over the 17 patients included in the present study. The set ℋ is a subset of *h* comprising the pressure and ejection fraction indices but not the volume indices used in the model parameter fitting. Due to the underestimation of volume measurements collected with TTE, the model fit of the volumes was evaluated by comparing the Pearson correlation coefficient between the simulated volume indices and clinical TTE volume measurements to the Pearson correlation coefficient between TTE and CMR imaging volume measurements reported in (Whitlock et al., 2010; Kühl et al., 2012). To assess whether the model can replicate hemodynamic changes between NSR and AF, a paired student’s t-test was applied to the simulated hemodynamic indices described in Section 2.4.2. A *p*-value < 0.05 was considered significant.

To evaluate the impact of different atrial contraction patterns on the model’s ability to replicate clinical measurements in AF, the absolute simulation error |*ϵ*_ℋ_| was compared between simulations with the proposed model (S1), simulations with no atrial contraction (S2), and simulations with regular and synchronized atrial contraction (S3). For S2 and S3, a new model was fitted using the approach described in Section 2.4.3, with identical ventricular activity but modified atrial activity. Notably, parameter fitting for S2 required a 50% increase in *σ*_*h*_ for all hemodynamic indices *h* listed in Table 4, as solutions could not otherwise be found for all patients. The ventricular activation times were given by ***ν***_*S*_ in 13 patients and by ***ν***_*C*_ in the remaining 4 patients with available ECG signals. For simulations with no atrial contraction (S2), none of the atrial wall patches were contracting throughout the simulation. For simulations with regular and synchronized atrial contraction (S3), all atrial wall patches were contracting at once 120 ms before each irregular ventricular activation in ***ν***_*S*_ or ***ν***_*C*_.

Furthermore, to illustrate the hemodynamic differences among the three atrial contraction patterns (S1, S2, and S3), a case study was conducted on patient P4, who had available EGM, ECG, and pressure measurements. For this patient, the simulated right atrial and right ventricular pressure trends of the fitted models S1, S2, and S3 were illustrated alongside clinical EGM, ECG, and pressure measurements.

## 3 Results

Section 3.1 focuses exclusively on evaluating the patient-specific fitting of the proposed model (S1), presenting individual fitting results for the 17 patients and summarizing simulation errors across the cohort. Section 3.2 then presents a direct comparison of patient-specific fits for three distinct atrial contraction patterns that describe the atrial activity in AF: the proposed model (S1), no atrial contraction (S2), and regular synchronized atrial contraction (S3).

### 3.1 Patient-specific model fitting

Figure 2 presents patient-specific model fits to arterial and intracardiac pressures for all 17 patients in both NSR and AF. Clinical measurements are shown alongside simulated pressures from models fitted using ***α***_*S*_ and ***ν***_*S*_, and, where available, models fitted using 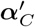 and ***ν***_*C*_. The simulated pressures were produced by each model using one final parameter set selected from 1024 parameter sets randomly drawn from the converged BHM parameter space as described in Section 2.4.3. Of these 1024 parameter sets, a median of 95.9% (range: 86.9%–100.0%) achieved computational convergence and were eligible for parameter selection. To illustrate prediction uncertainty, BHM posterior distributions are shown alongside each simulated pressure. These distributions, which were not part of parameter selection, include a median of 74.9% (range: 54.2%–81.1%) of the 1024 parameter sets with implausibility scores below 2. Comparison of model fits with and without ECG and EGM data shows a high similarity for 3 out of 4 patients (P4, P9, P15), suggesting that model fitting without ECG and EGM data is feasible in many cases. However, for patient P13, the model fitting using 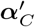 and ***ν***_*C*_ can match the clinical measurements of SBP, DBP, RVSP, and LAP_avg_ visibly better.

**Figure 2:**
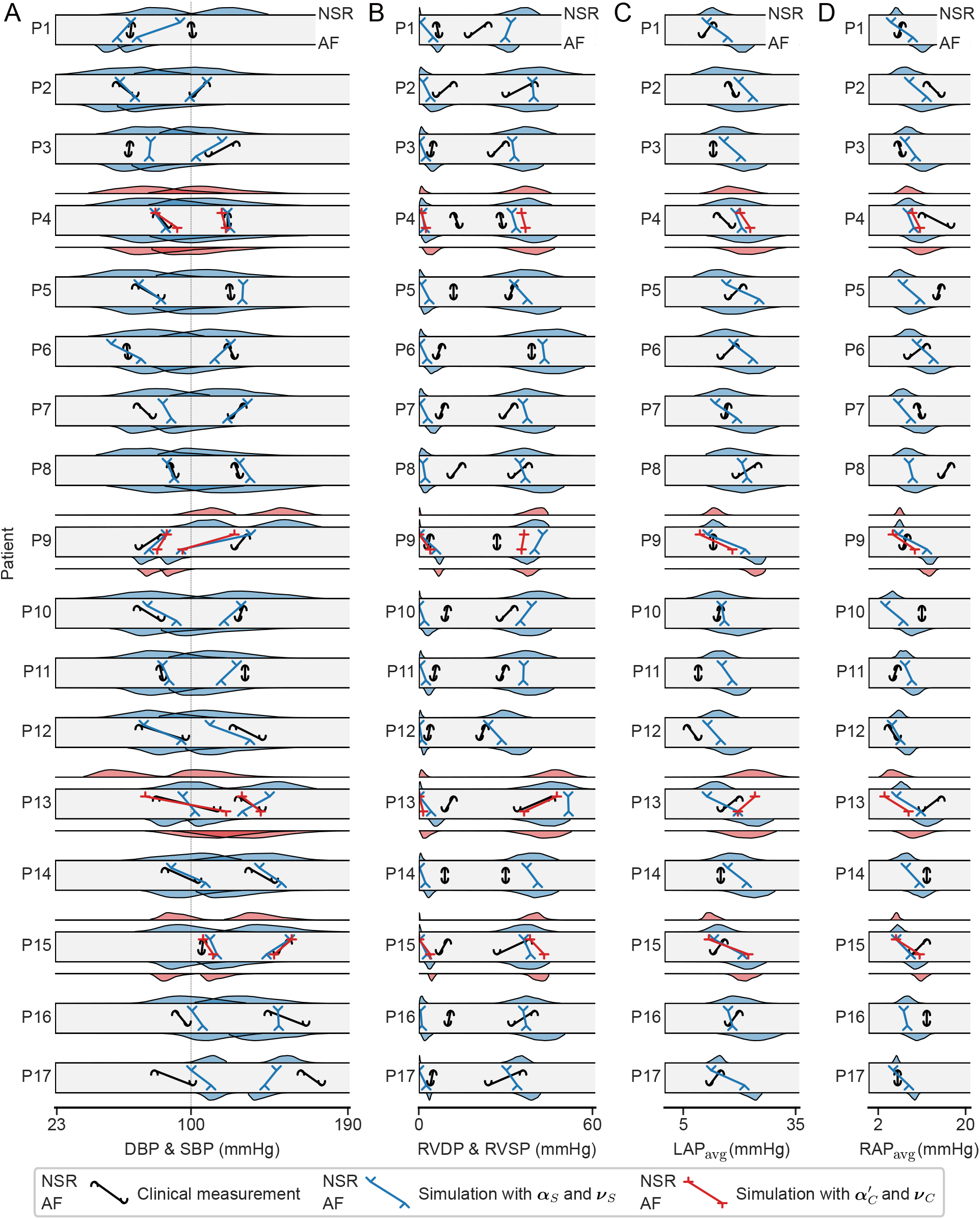
Results of model fitting, where each row (P1-P17) corresponds to one patient. **A)** diastolic and systolic blood pressure (DBP, SBP), **B)** right ventricular diastolic and systolic pressure (RVDP, RVSP), **C)** mean left atrial pressure (LAP_avg_), and **D)** mean right atrial pressure (RAP_avg_). The markers show the clinical measurements (black) and simulation results with the final parameter set, fitted using ***α***_*S*_ and ***ν***_*S*_ (blue) and using 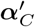 and ***ν***_*C*_ (red), if available, in (top) NSR and (bottom) AF. Along the top and bottom edge of a row, the posterior distribution of the Bayesian history matching is shown from which the final parameter set was selected with the cost function *J* defined in Eq. 6.

Table 5 summarizes the simulated hemodynamic indices of the patient-specific models in both NSR and AF, including arterial and intracardiac pressures, atrial and ventricular ejection fractions and volumes, and cardiac output. For hemodynamic indices in set ℋ, the normalized simulation error *ϵ*_*ℋ*,norm_ (Eq. 8) is also presented. In terms of model bias assessed via *ϵ*_*ℋ*,norm_, the model replicated most hemodynamic indices in set ℋ except for RVDP in NSR, which was consistently underestimated, and RVSP and LAP_avg_ in AF, which were consistently overestimated. Regarding hemodynamic indices without direct clinical measurements, comparison with reported physiological ranges indicated that atrial and ventricular ejection fractions (LAEF, RAEF, LVEF, RVEF) were accurately replicated within expected physiological ranges for both NSR and AF (Therkelsen et al., 2006; Aune et al., 2009; Tamborini et al., 2010; Kanagala et al., 2020). Notably, atrial ejection fraction decreased markedly in AF to approximately 4%, while ventricular ejection fraction decreased by approximately one-third, consistent with observations (Therkelsen et al., 2006). During NSR, the patient-specific models accurately reproduced cardiac volumes, with all volume indices (LAV_max_, RAV_max_, LVDV, RVDV) falling within sex-specific ranges reported by Fuchs et al. (2016). However, the models did not replicate the expected increase in cardiac volumes from NSR to AF (Therkelsen et al., 2006). Simulated atrial volumes demonstrated strong correlations with clinical TTE measurements (*r* = 0.76 for LAV_max_, *r* = 0.82 for RAV_max_), comparable with reported correlations between TTE and CMR imaging (*r* = 0.91 for LAV_max_ and 0.82 for RAV_max_ (Whitlock et al., 2010), *r* = 0.71 for LAV_max_ (Kühl et al., 2012)). Cardiac output remained within normal physiological ranges across both NSR and AF (Klavebäck et al., 2023). Although a mean reduction in cardiac output was observed from NSR to AF, consistent with previous observations (Klavebäck et al., 2023), this reduction did not achieve statistical significance. Table 6 summarizes the parameters of the fitted models.

**Table 5:** Simulation results of fitted model and normalized simulation error *ϵ*_*ℋ*,norm_ in NSR and AF. The errors *ϵ*_*ℋ*,norm_ were computed according to Eq. 8.

| Variables | Simulation result | | Normalized simulation error $\epsilon_{\mathcal{H},\text{norm}}$ | |
| --- | --- | --- | --- | --- |
|  | NSR | AF | NSR | AF |
| SBP (mmHg) | 127.8 [93.7, 157.8] | 120.9 [69.0, 152.0] | 0.01 [-0.93, 0.53] | -0.35 [-1.62, 0.32] |
| DBP (mmHg) | 79.6 [54.3, 106.9] | 90.3 [57.6, 120.1] | 0.17 [-0.70, 1.70] | 0.32 [-0.46, 0.74] |
| RVSP (mmHg) | 36.1 [24.0, 47.7] | 36.9 [28.7, 43.4] | 0.36 [-0.91, 1.50] | 1.60 [0.48, 3.25] |
| RVDP (mmHg) | 0.4 [0.1, 1.5] | 2.5 [1.1, 5.0]* | -2.60 [-4.07, -1.14] | -0.97 [-3.60, 0.30] |
| LAP<sub>avg</sub> (mmHg) | 15.8 [9.5, 24.1] | 20.4 [15.0, 25.3]* | -0.13 [-0.99, 1.38] | 1.89 [0.36, 2.94] |
| RAP<sub>avg</sub> (mmHg) | 7.1 [3.3, 9.9] | 9.1 [6.6, 13.4]* | -0.99 [-3.19, 0.41] | -0.4 [-1.72, 1.45] |
| LAEF (%) | 56.7 [44.0, 75.8] | 4.4 [1.3, 10.5]* | 0.54 [-1.03, 1.62] | — |
| RAEF (%) | 45.4 [37.3, 77.5] | 4.1 [2.5, 11.7]* | — | — |
| LVEF (%) | 61.4 [51.7, 71.6] | 42.2 [22.1, 55.7]* | -0.21 [-2.42, 1.31] | — |
| RVEF (%) | 56.3 [30.9, 84.0] | 31.0 [9.69, 61.5]* | — | — |
| LAV<sub>max</sub> (mL) | 75.0 [54.7, 111.1] | 35.2 [21.4, 66.3]* | — | — |
| RAV<sub>max</sub> (mL) | 84.4 [57.0, 135.4] | 43.9 [26.6, 74.6]* | — | — |
| LVDV (mL) | 125.2 [96.3, 169.7] | 104.0 [57.3, 135.2]* | — | — |
| RVDV (mL) | 144.9 [105.7, 179.4] | 124.9 [93.5, 178.4]* | — | — |
| CO (L·min<sup>-1</sup>) | 4.52 [2.69, 7.43] | 3.74 [1.43, 6.48] | — | — |

Hemodynamic indices and errors are presented as median [min, max]. CO, cardiac output; DBP, diastolic blood pressure; LAEF, left atrial ejection fraction; LAP<sub>avg</sub>, mean left atrial pressure; LAV<sub>max</sub>, maximum left atrial volume; LVDV, left ventricular diastolic volume; LVEF, left ventricular ejection fraction; RAEF, right atrial ejection fraction; RAP<sub>avg</sub>, mean right atrial pressure; RAV<sub>max</sub>, maximum right atrial volume; RVDP, right ventricular diastolic pressure; RVDV, right ventricular diastolic volume; RVEF, right ventricular ejection fraction; RVSP, right ventricular systolic pressure; SBP, systolic blood pressure. \* marks significant increase/decrease from NSR to AF according to a paired student's t-test ( $p < 0.05$ ).

**Table 6:** Fitted model parameters of the final parameter sets.

| Parameter | Estimate | Unit |
| --- | --- | --- |
| $A_{m,ref,RA}$ | 9096 [6353, 12086] | mm<sup>2</sup> |
| $A_{m,ref,LA}$ | 7222 [4968, 8291] | mm<sup>2</sup> |
| $A_{m,ref,RFW}$ | 16410 [12282, 21081] | mm<sup>2</sup> |
| $A_{m,ref,SW}$ | 6376 [3702, 7977] | mm<sup>2</sup> |
| $A_{m,ref,LFW}$ | 6259 [4404, 8376] | mm<sup>2</sup> |
| $S_{act,v}$ | 0.477 [0.160, 0.587] | MPa |
| $S_{act,a}$ | 0.387 [0.108, 0.600] | MPa |
| $t_{ad,v}$ | 318 [202, 394] | ms |
| $t_{ad,a}$ | 306 [88, 375] | ms |
| $k_1$ | 1.348 [1.071, 1.398] | – |
| $SBV$ | 1356 [1046, 1834] | mL |
| $R_{as}$ | 1.372 [0.585, 1.950] | mmHg·s·mL<sup>-1</sup> |
| $R_{ap}$ | 0.075 [0.026, 0.279] | mmHg·s·mL<sup>-1</sup> |
| $k_{Cs}$ | 0.926 [0.812, 1.130] | – |
| $k_{Cp}$ | 1.015 [0.806, 1.134] | – |
| $k_2$ | 1.208 [1.048, 1.399] | – |

The parameters of the fitted models are presented as median [min, max].

Figure 3A and 3B illustrate the magnitude of the simulation error *ϵ*_*ℋ*_ (Eq. 7) for the hemodynamic indices in ℋ during NSR and AF when using the proposed model (S1). Regarding model accuracy, 75% of absolute pressure simulation errors were below 8.1 mmHg, and 75% of absolute ejection fraction simulation errors were below 8.6%. The median of the average pressure simulation error across patients was 6.0 mmHg (range: 3.4−12.6 mmHg) for both NSR and AF. During NSR, the median of the average ejection fraction simulation error was 5.2% (range: 0.5−11.0%).

**Figure 3:**
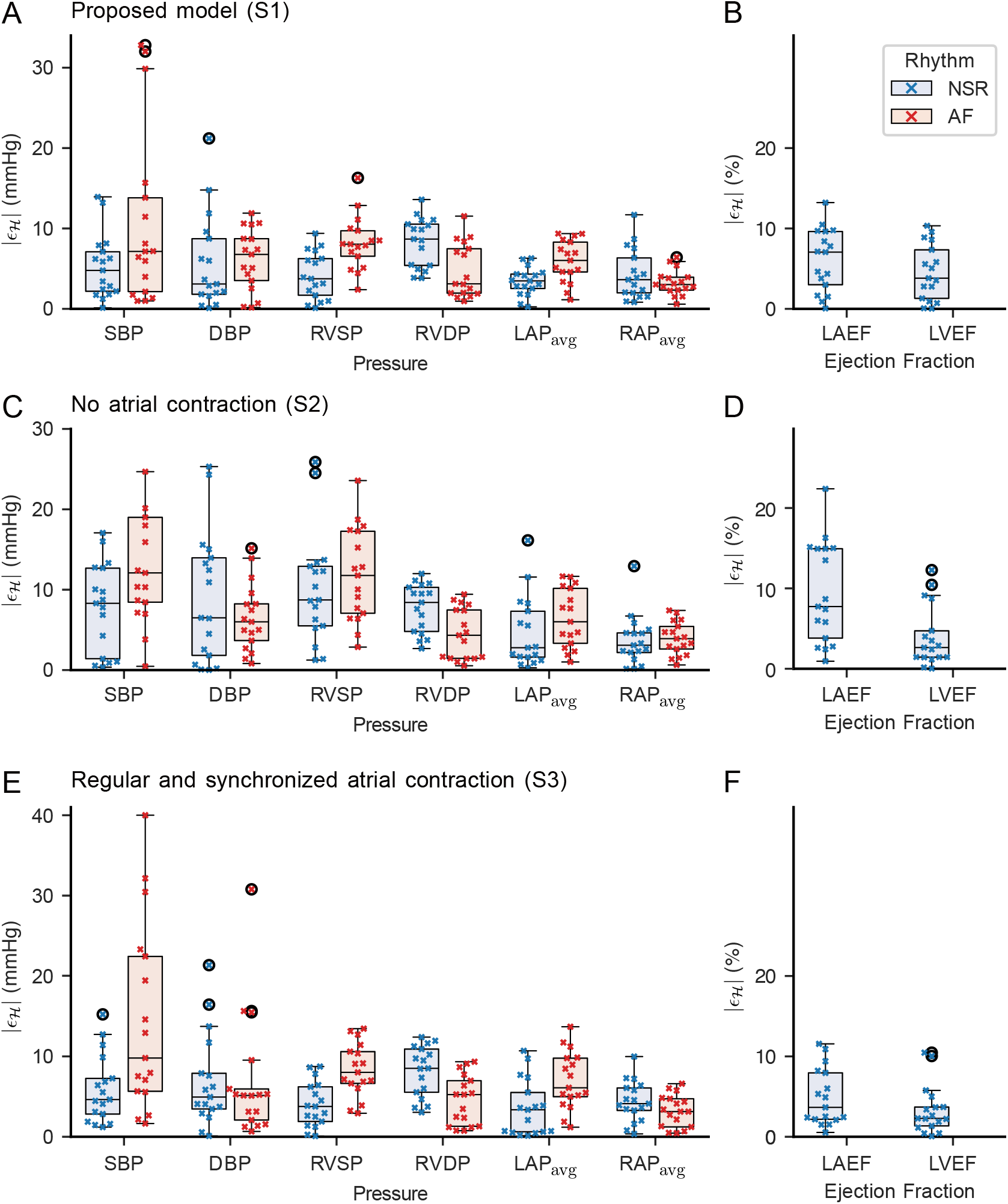
Magnitude of the simulation error *ϵ*_*ℋ*_ for hemodynamic indices during NSR and AF for simulations with the proposed model (S1, **A**+**B**), simulations with no atrial contraction (S2, **C**+**D**), and simulations with regular and synchronized atrial contraction (S3, **E**+**F**). **A**+**C**+**E**: Pressure simulation errors |*ϵ* _ℋ_| during NSR (left, blue) and AF (right, red). **B**+**D**+**F**: Ejection fraction simulation errors |*ϵ*_ℋ_| during NSR. Simulation errors of individual patients (crosses, n=17) are shown alongside boxplots summarizing the simulation errors of all 17 patients.

### 3.2 Comparison to other models

Figure 3 compares the absolute simulation error *ϵ*_*ℋ*_ for the hemodynamic indices in ℋ during NSR and AF across three atrial contraction patterns describing the atrial activity in AF: the proposed model (S1, A+B), no atrial contraction (S2, C+D), and regular and synchronized atrial contraction (S3, E+F). Compared to the proposed model (S1), models simulating no atrial contraction (S2) exhibited higher absolute simulation errors, where 75% of absolute pressure simulation errors were below 10.5 mmHg and 75% of absolute ejection fraction simulation errors were below 10.1%. The median of the average pressure simulation error for S2 was 8.2 mmHg (range: 5.4−15.8 mmHg), and the median of the average ejection fraction simulation error was 5.7% (range: 1.3−13.6%). Models simulating regular and synchronized atrial contraction during AF (S3) demonstrated mixed performance relative to the proposed model (S1), with higher absolute pressure simulation errors (75% below 8.5 mmHg) but lower absolute ejection fraction simulation errors (75% below 5.7%). The median of the average pressure simulation error for S3 was 6.6 mmHg (range: 5.2−15.5 mmHg), and the median of the average ejection fraction simulation error was 3.1% (range: 1.0−10.8%).

To further evaluate the simulated hemodynamics in the atria and ventricles as modeled with the three atrial contraction patterns (S1, S2, S3), a case study with patient P4 illustrates 10-second pressure trends in the right atrium (Figure 4) and right ventricle (Figure 5). Both figures compare simulated and clinical pressure trends for each atrial contraction pattern. Patient P4 was chosen for this case study, as they were the only patient with available clinical pressure trends. Notably, patient P4 exhibited the highest clinical RAP_avg_, RV systolic and diastolic pressure and represented for the proposed model the largest absolute difference between clinical and simulated RAP_avg_, RV systolic and diastolic pressure within the study cohort (Figure 2, Figure 3A). Despite offsets between simulated and clinical pressure trends, the case study illustrates key qualitative similarities between the simulated and clinical pressure trends.

**Figure 4:**
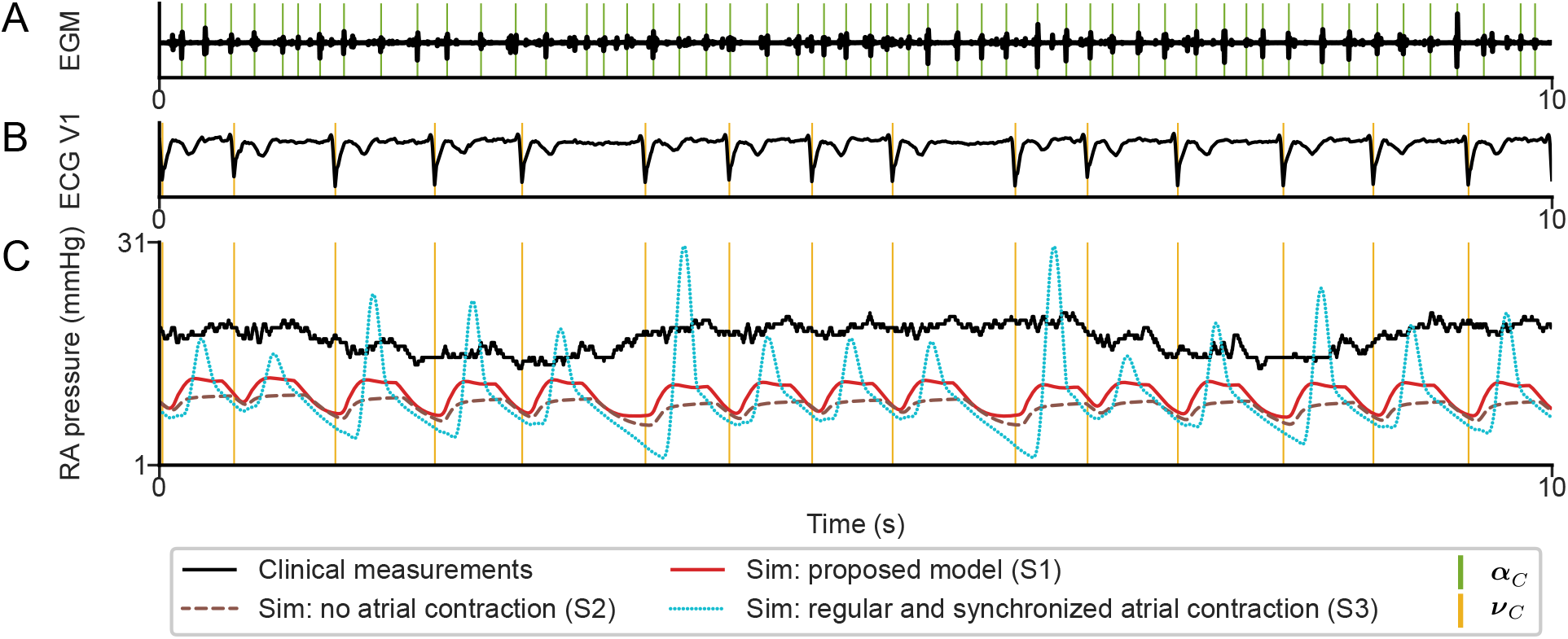
Right atrial (RA) pressure during atrial fibrillation. **A)** Intracardiac electrogram signal (black) from a catheter in the coronary sinus with the series of atrial activation times ***α***_*C*_ (green). **B)** ECG V1 lead (black) with the series of ventricular activation times ***ν***_*C*_ (orange). **C**) Right atrial pressure measured with catheter (black), simulated with the proposed model (S1, red), simulated with no atrial contraction (S2, brown), and simulated with regular and synchronized atrial contraction (S3, blue).

**Figure 5:**
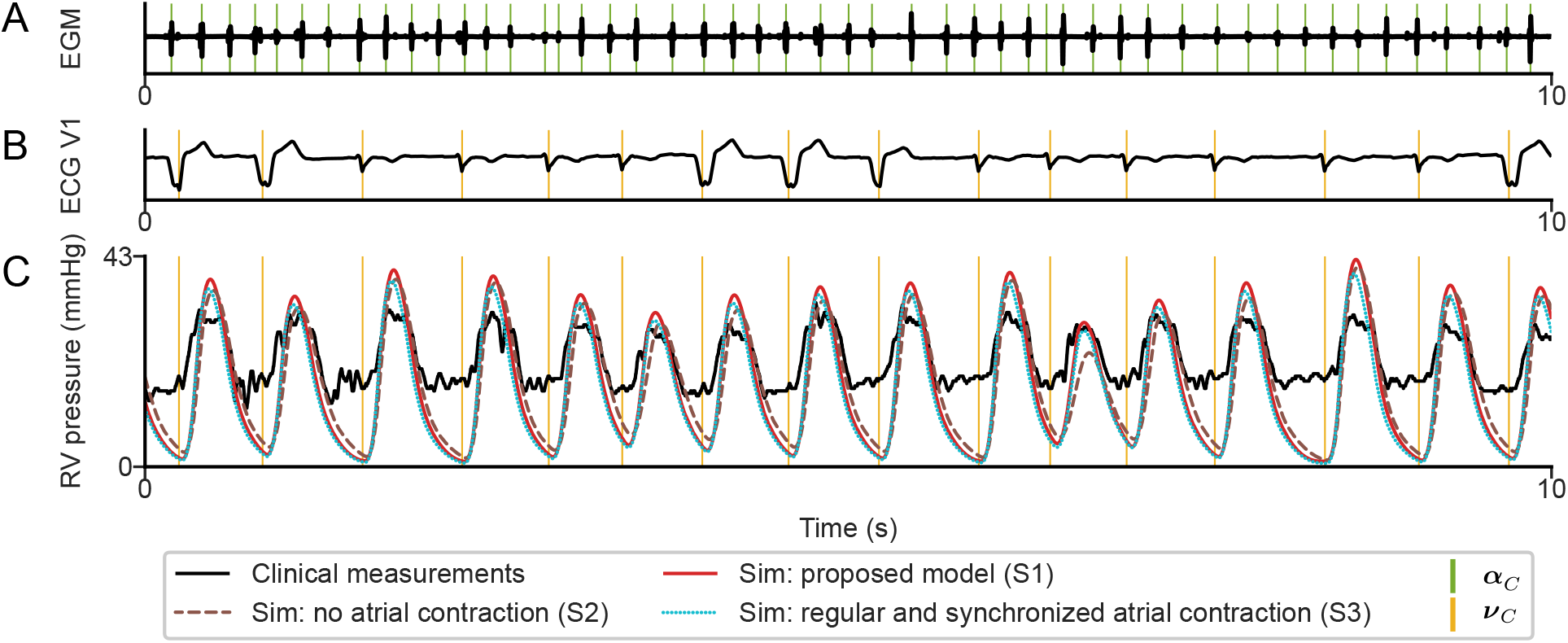
Right ventricular (RV) pressure during atrial fibrillation. **A)** Intracardiac electrogram signal (black) from a catheter in the coronary sinus with the series of atrial activation times ***α***_*C*_ (green). **B)** ECG V1 lead (black) with the series of ventricular activation times ***ν***_*C*_ (orange). **C**) Right ventricular pressure measured with catheter (black), simulated with the proposed model (S1, red), simulated with no atrial contraction (S2, brown), and simulated with regular and synchronized atrial contraction (S3, blue).

In Figure 4C, all three models (S1, S2, S3) underestimated right atrial pressure, yielding absolute simulation errors of 6.4 mmHg, 3.1 mmHg, and 6.0 mmHg, respectively. Both the proposed model (S1) and the model using no atrial contraction (S2) produced right atrial pressure variations of similar magnitude to the clinical measurement, whereas the model using a regular and synchronized atrial contraction (S3) produced atrial pressure variations within a cardiac cycle that were much larger than those of the clinical measurement. None of the models replicated the slow baseline variation in the clinical pressure trend that ranged multiple heartbeats, though beat-to-beat variation in simulated pressures relative to the preceding diastolic interval can be seen.

Simulated RV pressure trends in Figure 5C were nearly identical across the three models, indicating that ventricular pressure remained largely independent of the atrial contraction pattern in the computational model. The clinical RV pressure trends displayed blunted extrema, in contrast to the simulated signals, which exhibited more pronounced systolic and diastolic peaks. Simulated RV systolic pressures were 36.9 mmHg (S1), 39.0 mmHg (S2), and 32.2 mmHg (S3), all higher than the clinical measurement of 29.0 mmHg. Simulated RV diastolic pressure were underestimated at 2.5 mmHg (S1), 5.8 mmHg (S2), and 4.9 mmHg (S3), compared to the clinical measurement of 14.0 mmHg.

## 4 Discussion

This study presents a computationally efficient model for simulating hemodynamics in AF. The model comprises two interconnected subsystems: an electrical subsystem generating atrial and ventricular activation times characteristic of AF, and a mechanical subsystem describing cardiovascular mechanics and hemodynamics. The model was fitted to 17 patients from the SMURF study to reproduce arterial and intracardiac pressure measurements in NSR and AF and ejection fraction measurements in NSR (Charitakis et al., 2016). Results in Figure 3A show that the proposed model matched the pressure measurements (SBP, DBP, RVSP, RVDP, LAP_avg_, RAP_avg_) in both NSR and AF with an absolute simulation error below 8.1 mmHg in a large majority (75%). Furthermore, the proposed model matched ejection fraction measurements (LAEF, LVEF) in NSR in a large majority with an absolute simulation error below 8.6% (Figure 3B).

When compared to two models with different atrial contraction patterns in AF, the proposed model achieved the lowest pressure simulation errors while producing atrial pressure variations within a cardiac cycle consistent with clinical measurements. Although a model with regular and synchronized atrial contraction yielded lower absolute ejection fraction simulation errors, it produced much larger atrial pressure variations than the clinical measurement (Figure 4C). The computational complexity of the model allows the simulation of beat-to-beat variation in hemodynamics and can simulate 100 heartbeats on average in 1 second on just a single CPU core. The median model fitting time per patient was 46 min (34 to 89 min) using 10 cores on an Apple M1 Pro chip and 32 GB of RAM.

### 4.1 Model design

The TriSeg model, using the MultiPatch approach (Arts et al., 2005; Lumens et al., 2009; Walmsley et al., 2015), provides a suitable framework for continuous hemodynamic simulations over many heartbeats to capture the beat-to-beat variation in AF and is efficient enough to simulate many realizations to enable patient-specific model parameter fitting. A limitation of the TriSeg model is its reliance on external input for electrical activation times of each wall patch. Previous studies have determined the electrical activation times for the wall patches by electrocardiographic mapping (Lumens et al., 2013), simulation using a 3D finite-element model (Meiburg et al., 2023), assuming simultaneous contraction of wall patches (Lyon et al., 2021), assuming no contraction of wall patches to simulate atrial activity in AF (Lyon et al., 2021), or using a simplified description of the cardiac conduction system (Taconné et al., 2022). This work proposes a novel electrical subsystem that complements the MultiPatch implementation of the TriSeg model by generating activation times characteristic of AF for each atrial and ventricular wall patch without requiring clinical measurements, while preserving the option to personalize heart rhythms using available ECG signals. We considered simulating electrical excitation propagation in AF using a network model, either based on the work of Taconné et al. (2022) or by extending the AV node network model to connect with atrial and ventricular wall patches. However, to facilitate fast computation times and to keep the number of model parameters low for model fitting, a stochastic approach for electrical activation timing was implemented. This stochastic approach is well-suited to the MultiPatch implementation, which does not specify the shape, location or adjacency of the patches. Without spatial information in the 0-dimensional model, electrical excitation can be approximated by how much tissue with comparable properties is activated at a given time without specifying the pattern or path of electrical excitation propagation. This stochastic approach of electrical activation timing serves as a surrogate when cardiac activation maps are unavailable.

The proposed model reproduces hemodynamic differences between NSR and AF through three mechanisms. First, atrial and ventricular contraction duration (*t*_ad,a_ and *t*_ad,v_) were modified using scaling factor *k*_1_ to account for the documented shortening of cardiac ejection time in AF compared to NSR (Krohn and Magidson, 1966; Tavel et al., 1972). Second, vascular parameters (*R*_as_, *R*_ap_, *C*_as_, *C*_vs_, *C*_ap_, *C*_vp_) were modified using scaling factor *k*_2_ to account for the documented increase in vascular resistance and decrease in vascular capacitance in AF compared to NSR (Gosselink et al., 1996; Tuinenburg et al., 1998). Because the effect of increased sympathetic activity is mimicked by an increase in *k*_1_ (Stafford et al., 1970) and *k*_2_ (Wieling et al., 1998; Svec et al., 2021), the model fitting restricted *k*_1_ and *k*_2_ to be either both *≥*1 or both *<*1. Third, the series of atrial and ventricular electrical activation times (***α*** and ***ν***) differ between NSR and AF to account for the differences in electrical activation patterns. Specifically, the uncoordinated atrial activity in AF was modeled by dividing each atrium into 20 wall patches with independently generated activation times. The atrial activation times were randomly drawn from a distribution, analogous to previous work that modeled atrial activations entering the AV node during AF (Cohen et al., 1983; Lian et al., 2006; Corino et al., 2011; Masè et al., 2015; Wallman and Sandberg, 2018; Plappert et al., 2022). Coordinated electrical activity was modeled as simultaneous whole-chamber activation, a simplification adopted due to missing clinical cardiac activation maps and because a sensitivity analysis (data not shown) indicated that inter-patch activation delays were not among the most influential parameters for patient-specific fitting.

### 4.2 Patient-specific model fitting

To personalize the model, the midwall reference area *A*_m,ref_, active stress *S*_act_, contraction duration *t*_ad_, stressed blood volume *SBV*, vascular resistance *R*, and vascular capacitance *C* were fitted to each patient and remained fixed between NSR and AF. These parameters, listed in Table 3 with fitted values in Table 6, were selected based on a sensitivity analysis (data not shown). These or closely related parameters have been employed previously to fit the TriSeg model to experimental or clinical data (Koopsen et al., 2024; Jones and Oomen, 2025). To replicate the ejection fraction estimates between 41–71%, the atrial and ventricular active myofiber stress *S*_act,a_ and *S*_act,v_ had to be fitted to values up to 4 times larger than in previous work (Arts et al., 2005; Lumens et al., 2009; Oomen et al., 2022; Koopsen et al., 2024; Jones and Oomen, 2025). In addition, ANS-induced changes to contraction duration and vasculature were accounted for by the scaling factors *k*_1_ and *k*_2_. Although the parameters *k*_1_ and *k*_2_ were fitted between 0.6 and 1.4, the fitting resulted in *k*_1_ *>* 1 and *k*_2_ *>* 1 in all 17 patients, consistent with an increased sympathetic response during AF.

The model replicated most hemodynamic indices in set ℋ except for RVDP in NSR, which was consistently under-estimated, and RVSP and LAP_avg_ in AF, which were consistently overestimated (Table 5). For the hemodynamic indices that were replicated and had available measurements in NSR and AF, the model captured characteristic changes from NSR to AF, such as an increase in DBP (Maselli et al., 2016; Olbers et al., 2019, 2021; Almroth et al., 2023) and an increase in RAP_avg_ (Clark et al., 1997). For the hemodynamic measurements without measurements in both NSR and AF, the model captured characteristic changes from NSR to AF, such as a decrease in LAEF, right atrial ejection fraction (RAEF), right ventricular ejection fraction (RVEF) (Therkelsen et al., 2006), and a decrease in LVEF (Kieny et al., 1992; Van Gelder et al., 1993; Therkelsen et al., 2006). Although a mean reduction in cardiac output was observed from NSR to AF, consistent with previous observations (Klavebäck et al., 2023), this reduction did not achieve statistical significance. The models did not replicate the expected increase in cardiac volumes from NSR to AF (Therkelsen et al., 2006).

### 4.3 Comparison to other models

Figure 3 compares the absolute simulation error *ϵ*_*ℋ*_ for the hemodynamic indices in ℋ during NSR and AF across three atrial contraction patterns describing the atrial activity in AF: the proposed model (S1, A+B), no atrial contraction (S2, C+D), and regular and synchronized atrial contraction (S3, E+F). No atrial contraction (S2) was chosen for the comparison because no atrial contraction with irregular ventricular activity recorded in patients was previously used to simulate atrial fibrillation (Lyon et al., 2021). Regular and synchronized atrial contraction (S3) was chosen to evaluate the hemodynamic differences between synchronized and uncoordinated atrial activation. The proposed model (S1) achieved the lowest absolute pressure simulation errors among the three atrial contraction patterns. While the model with regular and synchronized atrial contraction (S3) showed slightly worse absolute pressure simulation errors, it better matched ejection fraction measurements than the proposed model (S1). The right atrial pressure trends (Figure 4) revealed that fitting atrial pressure using its mean value alone masked important differences between the models. Despite similar absolute pressure errors, the model with regular and synchronized atrial contraction (S3) produced pressure variations substantially larger than the clinical measurement, whereas the proposed model closely matched the atrial pressure variation. Therefore, it is concluded that the proposed model best replicates the hemodynamics in both NSR and AF. Furthermore, the model without atrial contraction during AF performed worst in matching the clinical data, demonstrating that some degree of atrial contractile function remains essential for simulating hemodynamics in AF.

### 4.4 Limitations

Several limitations of this study warrant discussion. While the presented model replicates known AF-induced changes by modifying atrial and ventricular contraction duration, vascular resistance and capacitance, and atrial and ventricular electrical activation timing, it does not incorporate all AF-related mechanisms. Notably, hemodynamic reflexes such as those mediated via the baroreceptor were not incorporated, which could significantly influence the simulated hemodynamics and potentially improve the model’s accuracy in predicting atrial and ventricular pressures and volumes (Sharifi et al., 2024). Furthermore, the presented model simulated coordinated electrical activity as whole-chamber activation. Although inter-patch activation delay was not among the most influential parameters for patient-specific fitting, incorporating a staggered cardiac excitation could further improve the patient-specific fitting. Additionally, the proposed model assumed identical mean atrial arrival rates *µ* across all atrial wall patches during AF. However, a simulation study showed that the atrial fibrillatory rate (AFR) corresponding to *µ* varies across the atria Sánchez et al. (2017). Another simplification was that the proposed model described NSR as perfectly regular series *RR*_*V*_ = 0), whereas NSR inherently exhibits variability (*RR*_*V*_ *≈* 0.08) (McManus et al., 2013).

## 5 Conclusions

This study presents a computationally-efficient model for simulating hemodynamics in atrial fibrillation. Together with the model fitting using bayesian history matching, the model allows for the study of patient-specific hemodynamics. By capturing beat-to-beat variations in cardiac activation, the model provides a computational framework for exploring hemodynamic responses during irregular heart rhythms. A stochastic approach for electrical activation timing was implemented to model the uncoordinated atrial activity during AF while facilitating fast computation times and keeping the number of model parameters low for model fitting. The results demonstrated that the proposed model best matched clinical measurements when compared to a model simulating no atrial contraction and a model simulating regular and synchronized atrial contraction in AF. The presented model that 1) includes a description of atrial and ventricular activation times characteristic of AF, 2) is able to perform continuous hemodynamic simulations over many heartbeats to capture the beat-to-beat variation, and 3) is efficient enough to simulate many realizations to enable patient-specific model parameter fitting, represents a valuable tool for studying the effects of heart rate and rhythm on hemodynamics during AF.

## Data availability statement

The data used for this study cannot be shared publicly with respect to participating subjects. Requests to access these datasets should be directed to. These methodologies and results will be made publicly available on GitHub upon publication of this paper at http://github.com/BeatLabUCI/.

## Ethics statement

The Regional Ethical Review Board of the Faculty of Health Sciences, Linköping, Sweden (registration number: 2011/40-31), approved the study protocol. All participants gave their written consent to participate in the study. This study complies with the Declaration of Helsinki.

## Author contributions

FP: Conceptualization, Data curation, Formal analysis, Investigation, Methodology, Project administration, Software, Validation, Visualization, Writing – original draft, Writing – review and editing. PO: Conceptualization, Formal analysis, Investigation, Methodology, Supervision, Writing – review and editing. CJ: Software, Writing – review and editing. EC: Data curation, Writing – review and editing. LK: Data curation, Writing – review and editing. PP: Conceptualization, Funding acquisition, Writing – review and editing. MW: Conceptualization, Formal analysis, Funding acquisition, Investigation, Methodology, Supervision, Validation, Writing – original draft, Writing – review and editing. FS: Conceptualization, Formal analysis, Funding acquisition, Investigation, Methodology, Project administration, Resources, Supervision, Validation, Writing – original draft, Writing – review and editing.

## Appendix

The target variance *σ*_*h*_ for pressure indices in the BHM was selected according to Eq. A1

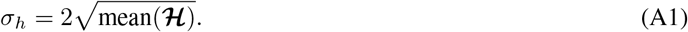

The pressure indices were divided into three groups: arterial pressures (SBP and DBP), right ventricular pressures (RVSP and RVDP), and atrial pressures (LAP_avg_ and RAP_avg_). For each group, **ℋ** contained measurements from all 17 patients during NSR and AF. This formulation was chosen to address the different pressure scales across the groups, providing an alternative to uniform or linearly proportional target variances that were producing either too small or too large target variances for some of the pressure indices.

The target variance *σ*_*h*_ for ejection fractions was selected separately due to different units from pressure indices. Since all ejection fractions were within the same order of magnitude, *σ*_*h*_ was uniformly set to 5. For volumes (LAV_max_, RAV_max_, LVDV, and RVDV), *µ*_*i*_ and *σ*_*i*_ were set to sex-specific means and standard deviations of atrial and ventricular volumes from Table 2 in (Fuchs et al., 2016).

| Parameter | Value | Unit | Source | Description |
| --- | --- | --- | --- | --- |
| $A_{m,ref,RA}$ | patient-specific [5 500, 12 500] | mm<sup>2</sup> | | RA midwall reference area |
| $A_{m,ref,LA}$ | patient-specific [4 500, 8 500] | mm<sup>2</sup> | | LA midwall reference area |
| $A_{m,ref,RFW}$ | patient-specific [11 000, 22 000] | mm<sup>2</sup> | | RFW midwall reference area |
| $A_{m,ref,SW}$ | patient-specific [3 500, 9 000] | mm<sup>2</sup> | | SW midwall reference area |
| $A_{m,ref,LFW}$ | patient-specific [4 000, 11 000] | mm<sup>2</sup> | | LFW midwall reference area |
| $RAW_{th}$ | 2.7 | mm | (Varela et al., 2017) | Right atrial wall thickness |
| $LAW_{th}$ | 2.4 | mm | (Varela et al., 2017) | Left atrial wall thickness |
| $RFW_{th}$ | 3.4 | mm | (Matsukubo et al., 1977) | Right ventricular free wall thickness |
| $SW_{th,1}$ | M:9.36, F:7.95 | mm | (Walpot et al., 2019) | Septal wall thickness Patch 2 |
| $SW_{th,2}$ | M:8.5, F:7.22 | mm | (Walpot et al., 2019) | Septal wall thickness Patch 3 |
| $SW_{th,3}$ | M:7.59, F:6.078 | mm | (Walpot et al., 2019) | Septal wall thickness Patch 8 |
| $SW_{th,4}$ | M:8.10, F:6.65 | mm | (Walpot et al., 2019) | Septal wall thickness Patch 9 |
| $SW_{th,5}$ | M:7.00, F:5.61 | mm | (Walpot et al., 2019) | Septal wall thickness Patch 14 |
| $LFW_{th,1}$ | M:9.21, F:7.47 | mm | (Walpot et al., 2019) | Left ventricular free wall thickness Patch 1 |
| $LFW_{th,2}$ | M:8.32, F:6.89 | mm | (Walpot et al., 2019) | Left ventricular free wall thickness Patch 4 |
| $LFW_{th,3}$ | M:8.68, F:7.03 | mm | (Walpot et al., 2019) | Left ventricular free wall thickness Patch 5 |
| $LFW_{th,4}$ | M:8.27, F:6.76 | mm | (Walpot et al., 2019) | Left ventricular free wall thickness Patch 6 |
| $LFW_{th,5}$ | M:7.27, F:5.95 | mm | (Walpot et al., 2019) | Left ventricular free wall thickness Patch 7 |
| $LFW_{th,6}$ | M:7.37, F:6.18 | mm | (Walpot et al., 2019) | Left ventricular free wall thickness Patch 10 |
| $LFW_{th,7}$ | M:7.56, F:6.01 | mm | (Walpot et al., 2019) | Left ventricular free wall thickness Patch 11 |
| $LFW_{th,8}$ | M:7.48, F:6.11 | mm | (Walpot et al., 2019) | Left ventricular free wall thickness Patch 12 |
| $LFW_{th,9}$ | M:7.53, F:6.51 | mm | (Walpot et al., 2019) | Left ventricular free wall thickness Patch 13 |
| $LFW_{th,10}$ | M:6.66, F:5.13 | mm | (Walpot et al., 2019) | Left ventricular free wall thickness Patch 15 |
| $LFW_{th,11}$ | M:7.46, F:6.05 | mm | (Walpot et al., 2019) | Left ventricular free wall thickness Patch 16 |

**Table A2:** Model parameters of the mechanical subsystem describing the cardiac mechanics and hemodynamics.

| Model | Parameter | Value | Unit | Source | Description |
| --- | --- | --- | --- | --- | --- |
| Constitutive<br>(active) | $L_{s,ref}$ | 2.0 | $\mu\text{m}$ | (Lumens et al., 2009) | Reference sarcomere length |
| | $L_{se,iso}$ | 0.04 | $\mu\text{m}$ | (Lumens et al., 2009) | Series elastic element length |
| | $L_{sc,0}$ | 1.51 | $\mu\text{m}$ | (Lumens et al., 2009) | Contractile element length at zero stress |
| | $v_{max,v}$ | 7 | $\mu\text{m} \cdot \text{s}^{-1}$ | (Lumens et al., 2009) | Maximum ventricular sarcomere shortening velocity |
| | $v_{max,a}$ | 14 | $\mu\text{m} \cdot \text{s}^{-1}$ | (Moyer et al., 2015) | Maximum atrial sarcomere shortening velocity |
| | $S_{act,v}$ | patient-specific [0.1, 0.6] | MPa | | Ventricular active stress coefficient |
| | $S_{act,a}$ | patient-specific [0.1, 0.6] | MPa | | Atrial active stress coefficient |
| | $\tau_{R,v}$ | $0.37 \cdot t_{ad,v}$ | ms | (Nollet et al., 2020) | Factor scaling ventricular contraction rise time |
| | $\tau_{D,v}$ | $0.37 \cdot t_{ad,v}$ | ms | (Nollet et al., 2020) | Factor scaling ventricular contraction decay time |
| | $\tau_{R,a}$ | $0.55 \cdot t_{ad,a}$ | ms | (Nollet et al., 2020) | Factor scaling atrial contraction rise time |
| | $\tau_{D,a}$ | $0.55 \cdot t_{ad,a}$ | ms | (Nollet et al., 2020) | Factor scaling atrial contraction decay time |
| | $t_{ad,v}$ | patient-specific [75, 500] $\cdot k_1^{-1}$ | ms | | Ventricular contraction duration |
| | $t_{ad,a}$ | patient-specific [75, 500] $\cdot k_1^{-1}$ | ms | | Atrial contraction duration |
| | $k_1$ | In NSR: 1<br>In AF: patient-specific [0.6, 1.4] | – | | Factor for changes in $t_{ad}$ between NSR and AF |
| Constitutive<br>(passive) | $c_{1,v}$ | 1e-5 | MPa | | Ventricular passive stress coefficient |
| | $c_{3,v}$ | 5.23e-4 | MPa | | Ventricular passive stress coefficient |
| | $c_{4,v}$ | 10 | – | | Ventricular passive stress coefficient |
| | $c_{1,a}$ | 1.60e-3 | MPa | | Atrial passive stress coefficient |
| | $c_{3,a}$ | 0.017e-3 | MPa | | Atrial passive stress coefficient |
| | $c_{4,a}$ | 10 | – | | Atrial passive stress coefficient |

**Table A3:**
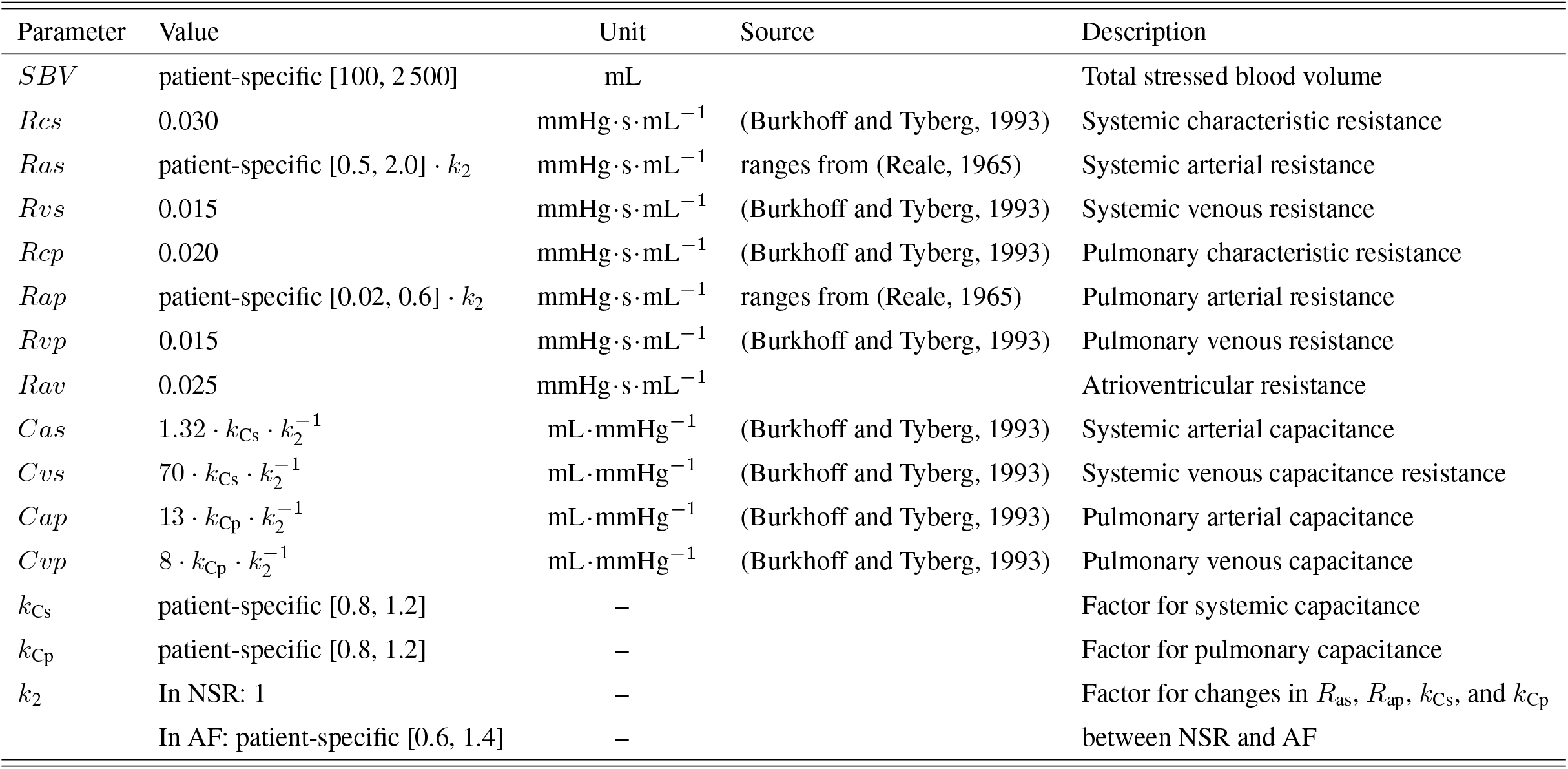
Model parameters of the mechanical subsystem describing the circulation system.

| Parameter | Value | Unit | Source | Description |
| --- | --- | --- | --- | --- |
| $SBV$ | patient-specific [100, 2 500] | mL | | Total stressed blood volume |
| $R_{cs}$ | 0.030 | mmHg·s·mL<sup>-1</sup> | (Burkhoff and Tyberg, 1993) | Systemic characteristic resistance |
| $R_{as}$ | patient-specific [0.5, 2.0] · $k_2$ | mmHg·s·mL<sup>-1</sup> | ranges from (Reale, 1965) | Systemic arterial resistance |
| $R_{vs}$ | 0.015 | mmHg·s·mL<sup>-1</sup> | (Burkhoff and Tyberg, 1993) | Systemic venous resistance |
| $R_{cp}$ | 0.020 | mmHg·s·mL<sup>-1</sup> | (Burkhoff and Tyberg, 1993) | Pulmonary characteristic resistance |
| $R_{ap}$ | patient-specific [0.02, 0.6] · $k_2$ | mmHg·s·mL<sup>-1</sup> | ranges from (Reale, 1965) | Pulmonary arterial resistance |
| $R_{vp}$ | 0.015 | mmHg·s·mL<sup>-1</sup> | (Burkhoff and Tyberg, 1993) | Pulmonary venous resistance |
| $R_{av}$ | 0.025 | mmHg·s·mL<sup>-1</sup> | | Atrioventricular resistance |
| $C_{as}$ | $1.32 \cdot k_{Cs} \cdot k_2^{-1}$ | mL·mmHg<sup>-1</sup> | (Burkhoff and Tyberg, 1993) | Systemic arterial capacitance |
| $C_{vs}$ | $70 \cdot k_{Cs} \cdot k_2^{-1}$ | mL·mmHg<sup>-1</sup> | (Burkhoff and Tyberg, 1993) | Systemic venous capacitance resistance |
| $C_{ap}$ | $13 \cdot k_{Cp} \cdot k_2^{-1}$ | mL·mmHg<sup>-1</sup> | (Burkhoff and Tyberg, 1993) | Pulmonary arterial capacitance |
| $C_{vp}$ | $8 \cdot k_{Cp} \cdot k_2^{-1}$ | mL·mmHg<sup>-1</sup> | (Burkhoff and Tyberg, 1993) | Pulmonary venous capacitance |
| $k_{Cs}$ | patient-specific [0.8, 1.2] | – | | Factor for systemic capacitance |
| $k_{Cp}$ | patient-specific [0.8, 1.2] | – | | Factor for pulmonary capacitance |
| $k_2$ | In NSR: 1<br>In AF: patient-specific [0.6, 1.4] | – | | Factor for changes in $R_{as}$, $R_{ap}$, $k_{Cs}$, and $k_{Cp}$ between NSR and AF |

## References

[1] Gerhard Hindricks, Tatjana Potpara, Nikolaos Dagres, Elena Arbelo, Jeroen J Bax, Carina Blomström-Lundqvist, Giuseppe Boriani, Manuel Castella, Gheorghe-Andrei Dan, Polychronis E Dilaveris, et al. 2020 ESC Guidelines for the diagnosis and management of atrial fibrillation developed in collaboration with the European Association of Cardio-Thoracic Surgery (EACTS). European Heart Journal, 42(5):373–498, 2020.

[2] James J. Morris, Mark Entman, William C. North, Yihong Kong, and Henry McIntosh. The Changes in Cardiac Output with Reversion of Atrial Fibrillation to Sinus Rhythm. Circulation, 31(5):670–678, 1965.

[3] Emile G. Daoud, Raul Weiss, Marwan Bahu, Bradley P. Knight, Frank Bogun, Rajiva Goyal, Mark Harvey, S. Adam Strickberger, K. Ching Man, and Fred Morady. Effect of an irregular ventricular rhythm on cardiac output. The American Journal of Cardiology, 78(12):1433–1436, 1996.

[4] Sofia Klavebäck, Helga Skúladóttir, Joakim Olbers, Jan Östergren, and Frieder Braunschweig. Changes in cardiac output, rhythm regularity, and symptom severity after electrical cardioversion of atrial fibrillation. Scandinavian Cardiovascular Journal, 57(1):2236341, 2023.

[5] Marta Carrara, Luca Carozzi, Travis J Moss, Marco de Pasquale, Sergio Cerutti, Manuela Ferrario, Douglas E Lake, and J Randall Moorman. Heart rate dynamics distinguish among atrial fibrillation, normal sinus rhythm and sinus rhythm with frequent ectopy. Physiological Measurement, 36(9):1873–1888, 2015.

[6] Rayo Akande, James Brimicombe, Martin R Cowie, Andrew Dymond, Hannah Clair Lindén, Gregory Y H Lip, Jenny Lund, Jonathan Mant, Madhumitha Pandiaraja, Emma Svennberg, et al. Characterising RR Intervals in Atrial Fibrillation Detected Through Screening. 2023 Computing in Cardiology (CinC), 50:1–4, 2023.

[7] Mamoru Kumada, Takehiko Azuma, and Kojiro Matsuda. The cardiac output-heart rate relationship under different conditions. The Japanese Journal of Physiology, 17(5):538, 1967.

[8] Joseph C. Greenfield, Alexander Harley, Howard K. Thompson, and Andrew G. Wallace. Pressure-flow studies in man during atrial fibrillation. Journal of Clinical Investigation, 47(10):2411–2421, 1968.

[9] Walter H Herbert. Cardiac Output and the Varying R-R Interval of Atrial Fibrillation. Journal of Electrocardiology, 6:131–135, 1973.

[10] Masahito Naito, Daniel David, Eric L. Michelson, Mark Schaffenburg, and Leonard S. Dreifus. The hemodynamic consequences of cardiac arrhythmias: Evaluation of the relative roles of abnormal atrioventricular sequencing, irregularity of ventricular rhythm and atrial fibrillation in a canine model. American Heart Journal, 106:284–291, 1983.

[11] David M Clark, Vance J Plumb, Andrew E Epstein, and G Neal Kay. Hemodynamic effects of an irregular sequence of ventricular cycle lengths during atrial fibrillation. J Am Coll Cardiol, 30:1039–1045, 1997.

[12] Isabelle C Van Gelder, Michiel Rienstra, Karina V Bunting, Ruben Casado-Arroyo, Valeria Caso, Harry J G M Crijns, Tom J R De Potter, Jeremy Dwight, Luigina Guasti, Thorsten Hanke, et al. 2024 ESC Guidelines for the management of atrial fibrillation developed in collaboration with the European Association for Cardio-Thoracic Surgery (EACTS): Developed by the task force for the management of atrial fibrillation of the European Society of Cardiology (ESC), with the special contribution of the European Heart Rhythm Association (EHRA) of the ESC. Endorsed by the European Stroke Organisation (ESO). European Heart Journal, 36(45):3314–3414, 2024.

[13] José A Joglar, Mina K Chung, Anastasia L Armbruster, Emelia J Benjamin, Janice Y Chyou, Edmond M Cronin, Anita Deswal, Lee L Eckhardt, Zachary D Goldberger, Rakesh Gopinathannair, et al. 2023 ACC/AHA/ACCP/HRS Guideline for the Diagnosis and Management of Atrial Fibrillation A Report of the American College of Cardiology/American Heart Association JointăCommitteeăonăClinical Practice Guidelines. Journal of the American College of Cardiology, 83(1):109–279, 2024.

[14] N.Sheldon Skinner, Jere H. Mitchell, Andrew G. Wallace, and Stanley J. Sarnoff. Hemodynamic consequences of atrial fibrillation at constant ventricular rates. The American Journal of Medicine, 36(3):342–350, 1964.

[15] Jere H. Mitchell and William Shapiro. Atrial function and the hemodynamic consequences of atrial fibrillation in man. The American Journal of Cardiology, 23(4):556–567, 1969.

[16] Charles B. Jr Upshaw. Hemodynamic changes after cardioversion of chronic atrial fibrillation. Archives of Internal Medicine, 157(10):1070–1076, 1997.

[17] Roy C. P. Kerckhoffs, Maxwell L. Neal, Quan Gu, James B. Bassingthwaighte, Jeff H. Omens, and Andrew D. McCulloch. Coupling of a 3D Finite Element Model of Cardiac Ventricular Mechanics to Lumped Systems Models of the Systemic and Pulmonic Circulation. Annals of Biomedical Engineering, 35(1):1–18, 2007.

[18] Christian B. Moyer, Patrick T. Norton, John D. Ferguson, and Jeffrey W. Holmes. Changes in Global and Regional Mechanics Due to Atrial Fibrillation: Insights from a Coupled Finite-Element and Circulation Model. Annals of Biomedical Engineering, 43(7):1600–1613, 2015.

[19] Marc Hirschvogel, Marina Bassilious, Lasse Jagschies, Stephen M. Wildhirt, and Michael W. Gee. A monolithic 3D0D coupled closedloop model of the heart and the vascular system: Experimentbased parameter estimation for patientspecific cardiac mechanics. International Journal for Numerical Methods in Biomedical Engineering, 33(8):e2842, 2017.

[20] Christoph M. Augustin, Matthias A.F. Gsell, Elias Karabelas, Erik Willemen, Frits W. Prinzen, Joost Lumens, Edward J. Vigmond, and Gernot Plank. A computationally efficient physiologically comprehensive 3D0D closed-loop model of the heart and circulation. Computer Methods in Applied Mechanics and Engineering, 386:114092, 2021.

[21] Tobias Gerach, Steffen Schuler, Jonathan Fröhlich, Laura Lindner, Ekaterina Kovacheva, Robin Moss, Eike Moritz Wülfers, Gunnar Seemann, Christian Wieners, and Axel Loewe. Electro-Mechanical Whole-Heart Digital Twins: A Fully Coupled Multi-Physics Approach. Mathematics, 9(11):1247, 2021.

[22] Serdar Göktepe, Jonathan Wong, and Ellen Kuhl. Atrial and ventricular fibrillation: computational simulation of spiral waves in cardiac tissue. Archive of Applied Mechanics, 80(5):569–580, 2010.

[23] Kensuke Sakata, Ryan P. Bradley, Adityo Prakosa, Carolyna A. P. Yamamoto, Syed Yusuf Ali, Shane Loeffler, Brock M. Tice, Patrick M. Boyle, Eugene G. Kholmovski, Ritu Yadav, et al. Assessing the arrhythmogenic propensity of fibrotic substrate using digital twins to inform a mechanisms-based atrial fibrillation ablation strategy. Nature Cardiovascular Research, 3(7):857–868, 2024.

[24] Joost Lumens, Tammo Delhaas, Borut Kirn, and Theo Arts. Three-Wall Segment (TriSeg) Model Describing Mechanics and Hemodynamics of Ventricular Interaction. Annals of Biomedical Engineering, 37:2234–2255, 2009.

[25] John Walmsley, Theo Arts, Nicolas Derval, Pierre Bordachar, Hubert Cochet, Sylvain Ploux, Frits W. Prinzen, Tammo Delhaas, and Joost Lumens. Fast Simulation of Mechanical Heterogeneity in the Electrically Asynchronous Heart Using the MultiPatch Module. PLoS Computational Biology, 11:1–23, 2015.

[26] Clara E. Jones and Pim J.A. Oomen. Synergistic biophysics and machine learning modeling to rapidly predict cardiac growth probability. Computers in Biology and Medicine, 184:109323, 2025.

[27] Pim J.A. Oomen, Thien Khoi N. Phung, Seth H. Weinberg, Kenneth C. Bilchick, and Jeffrey W. Holmes. A rapid electromechanical model to predict reverse remodeling following cardiac resynchronization therapy. Biomechanics and Modeling in Mechanobiology, 21:231–247, 2022.

[28] E Charitakis, U Walfridsson, F Nyström, E Nylander, A Strömberg, U Alehagen, and H Walfridsson. Symptom burden, Metabolic profile, Ultrasound findings, Rhythm, neurohormonal activation, haemodynamics and health-related quality of life in patients with atrial Fibrillation (SMURF): a protocol for an observational study with a randomised interventional component. BMJ Open, 5(12):e008723, 2015.

[29] Emmanouil Charitakis, Håkan Walfridsson, Eva Nylander, and Urban Alehagen. Neurohormonal Activation After Atrial Fibrillation Initiation in Patients Eligible for Catheter Ablation: A Randomized Controlled Study. Journal of the American Heart Association, 5(12):e003957, 2016.

[30] Henrik Almroth, Lars O Karlsson, Carl-Johan Carlhäll, and Emmanouil Charitakis. Haemodynamic changes after atrial fibrillation initiation in patients eligible for catheter ablation: a randomized controlled study. European Heart Journal Open, 3(6):oead112, 2023.

[31] Nicolas Pilia, Claudia Nagel, Gustavo Lenis, Silvia Becker, Olaf Dössel, and Axel Loewe. ECGdeliă-An open source ECG delineation toolbox for MATLAB. SoftwareX, 13:100639, 2021.

[32] Alejandro Alcaine, Michela Mase, Alessandro Cristoforetti, Flavia Ravelli, Giandomenico Nollo, Pablo Laguna, Juan Pablo Martinez, and Luca Faes. A Multi-Variate Predictability Framework to Assess Invasive Cardiac Activity and Interactions During Atrial Fibrillation. IEEE Transactions on Biomedical Engineering, 64(5):1157–1168, 2017.

[33] Steven M. Bishop and Ari Ercole. Multi-Scale Peak and Trough Detection Optimised for Periodic and Quasi-Periodic Neuroscience Data. In Intracranial Pressure {\&} Neuromonitoring XVI, volume 126, pages 189–195, 2018.

[34] Mattias Karlsson, Frida Sandberg, Sara R. Ulimoen, and Mikael Wallman. Non-invasive Characterization of Human AV-Nodal Conduction Delay and Refractory Period During Atrial Fibrillation. Frontiers in Physiology, 12, 2021.

[35] Theo Arts, Tammo Delhaas, Peter Bovendeerd, Xander Verbeek, and Frits W Prinzen. Adaptation to mechanical load determines shape and properties of heart and circulation: the CircAdapt model. Am J Physiol Heart Circ Physiol, 288:1943–1954, 2005.

[36] Andreu M. Climent, Felipe Atienza, Jose Millet, and Maria S. Guillem. Generation of realistic atrial to atrial interval series during atrial fibrillation. Medical and Biological Engineering and Computing, 49:1261–1268, 2011.

[37] Felix Plappert, Mikael Wallman, Mostafa Abdollahpur, Pyotr G. Platonov, Sten Östenson, and Frida Sandberg. An atrioventricular node model incorporating autonomic tone. Frontiers in Physiology, 13:976468, 2022.

[38] Mikael Wallman and Frida Sandberg. Characterisation of human AV-nodal properties using a network model. Medical and Biological Engineering and Computing, 56:247–259, 2018.

[39] W. P. Santamore and D. Burkhoff. Hemodynamic consequences of ventricular interaction as assessed by model analysis. American Journal of Physiology-Heart and Circulatory Physiology, 260(1):H146–H157, 1991.

[40] Nelson B. Schiller, Pravin M. Shah, Michael Crawford, Anthony DeMaria, Richard Devereux, Harvey Feigenbaum, Howard Gutgesell, Nathaniel Reichek, David Sahn, Ingela Schnittger, Norman H. Silverman, and A. Jamil Tajik. Recommendations for Quantitation of the Left Ventricle by Two-Dimensional Echocardiography. Journal of the American Society of Echocardiography, 2(5):358–367, 1989.

[41] David D. McManus, Jinseok Lee, Oscar Maitas, Nada Esa, Rahul Pidikiti, Alex Carlucci, Josephine Harrington, Eric Mick, and Ki H. Chon. A novel application for the detection of an irregular pulse using an iPhone 4S in patients with atrial fibrillation. Heart Rhythm, 10(3):315–319, 2013.

[42] Valentina D.A. Corino, Sara R. Ulimoen, Steve Enger, Luca T. Mainardi, Arnljot Tveit, and Pyotr G. Platonov. Rate-control drugs affect variability and irregularity measures of RR intervals in patients with permanent atrial fibrillation. Journal of Cardiovascular Electrophysiology, 26:137–141, 2015.

[43] I.M Sobol’. On the distribution of points in a cube and the approximate evaluation of integrals. USSR Computational Mathematics and Mathematical Physics, 7(4):86–112, 1967.

[44] Stephen Joe and Frances Y. Kuo. Constructing sobol sequences with better two-dimensional projections. SIAM Journal on Scientific Computing, 30(5):2635–2654, 2008.

[45] Edgar E. Nollet, Emmy M. Manders, Max Goebel, Valentijn Jansen, Cord Brockmann, Jorrit Osinga, Jolanda van der Velden, Michiel Helmes, and Diederik W. D. Kuster. Large-Scale Contractility Measurements Reveal Large Atrioventricular and Subtle Interventricular Differences in Cultured Unloaded Rat Cardiomyocytes. Frontiers in Physiology, 11:815, 2020.

[46] Ian Vernon, Michael Goldstein, and Richard G. Bower. Galaxy formation: a Bayesian uncertainty analysis. Bayesian Analysis, 5(4):619–669, 2010.

[47] Peter S. Craig, Michael Goldstein, Allan H. Seheult, and James A. Smith. Pressure Matching for Hydrocarbon Reservoirs: A Case Study in the Use of Bayes Linear Strategies for Large Computer Experiments. In Case Studies in Bayesian Statistics, Case Studies in Bayesian Statistics, pages 37–93. Springer New York, 1997.

[48] Matthew Whitlock, Anuj Garg, Jill Gelow, Timothy Jacobson, and Craig Broberg. Comparison of Left and Right Atrial Volume by Echocardiography Versus Cardiac Magnetic Resonance Imaging Using the Area-Length Method. The American Journal of Cardiology, 106(9):1345–1350, 2010.

[49] J. Tobias Kühl, Jacob Lønborg, Andreas Fuchs, Mads J. Andersen, Niels Vejlstrup, Henning Kelbæk, Thomas Engstrøm, Jacob E. Møller, and Klaus F. Kofoed. Assessment of left atrial volume and function: a comparative study between echocardiography, magnetic resonance imaging and multi slice computed tomography. The International Journal of Cardiovascular Imaging, 28(5):1061–1071, 2012.

[50] Tetsuji Kitano, Yosuke Nabeshima, Yutaka Otsuji, Kazuaki Negishi, and Masaaki Takeuchi. Accuracy of Left Ventricular Volumes and Ejection Fraction Measurements by Contemporary Three-Dimensional Echocardiography with Semi- and Fully Automated Software: Systematic Review and Meta-Analysis of 1,881 Subjects. Journal of the American Society of Echocardiography, 32(9):1105–1115.e5, 2019.

[51] Andreas Fuchs, Mads R. Mejdahl, J. Tobias Kühl, Zara R. Stisen, Emma Julia P. Nilsson, Lars V. Køber, Børge G. Nordestgaard, and Klaus F. Kofoed. Normal values of left ventricular mass and cardiac chamber volumes assessed by 320-detector computed tomography angiography in the Copenhagen General Population Study. European Heart Journal - Cardiovascular Imaging, 17(9):1009–1017, 2016.

[52] Sam Coveney and Richard H. Clayton. Fitting two human atrial cell models to experimental data using Bayesian history matching. Progress in Biophysics and Molecular Biology, 139:43–58, 2018.

[53] Susette Krohn Therkelsen, Bjoern Aaris Groenning, Jesper Hastrup Svendsen, and Gorm Boje Jensen. Atrial and Ventricular Volume and Function Evaluated by Magnetic Resonance Imaging in Patients With Persistent Atrial Fibrillation Before and After Cardioversion. The American Journal of Cardiology, 97(8):1213–1219, 2006.

[54] Erlend Aune, Morten Baekkevar, Jo Roislien, Olaf Rodevand, and Jan Erik Otterstad. Normal reference ranges for left and right atrial volume indexes and ejection fractions obtained with real-time three-dimensional echocardiography. European Journal of Echocardiography, 10(6):738–744, 2009.

[55] Gloria Tamborini, Nina Ajmone Marsan, Paola Gripari, Francesco Maffessanti, Denise Brusoni, Manuela Muratori, Enrico G. Caiani, Cesare Fiorentini, and Mauro Pepi. Reference Values for Right Ventricular Volumes and Ejection Fraction With Real-Time Three-Dimensional Echocardiography: Evaluation in a Large Series of Normal Subjects. Journal of the American Society of Echocardiography, 23(2):109–115, 2010.

[56] Prathap Kanagala, Jayanth R. Arnold, Adrian S. H. Cheng, Anvesha Singh, Jamal N. Khan, Gaurav S. Gulsin, Jing Yang, Lei Zhao, Pankaj Gupta, Iain B. Squire, et al. Left atrial ejection fraction and outcomes in heart failure with preserved ejection fraction. The International Journal of Cardiovascular Imaging, 36(1):101–110, 2020.

[57] Joost Lumens, Sylvain Ploux, Marc Strik, John Gorcsan, Hubert Cochet, Nicolas Derval, Maria Strom, Charu Ramanathan, Philippe Ritter, Michel Haïssaguerre, et al. Comparative Electromechanical and Hemodynamic Effects of Left Ventricular and Biventricular Pacing in Dyssynchronous Heart Failure Electrical Resynchronization Versus LeftRight Ventricular Interaction. Journal of the American College of Cardiology, 62(25):2395–2403, 2013.

[58] Roel Meiburg, Jesse H J Rijks, Ahmed S Beela, Edoardo Bressi, Domenico Grieco, Tammo Delhaas, Justin G LM Luermans, Frits W Prinzen, Kevin Vernooy, and Joost Lumens. Comparison of novel ventricular pacing strategies using an electro-mechanical simulation platform. Europace, 25(6):euad144, 2023.

[59] Aurore Lyon, Manouk van Mourik, Laura Cruts, Jordi Heijman, Sebastiaan C A M Bekkers, Ulrich Schotten, Harry J G M Crijns, Dominik Linz, and Joost Lumens. Both beat-to-beat changes in RR-interval and left ventricular filling time determine ventricular function during atrial fibrillation. EP Europace, 23(Supplement_1):i21–i28, 2021.

[60] Marion Taconné, Kimi P. Owashi, Elena Galli, Jürgen Duchenne, Arnaud Hubert, Erwan Donal, Alfredo I. Hernàndez, and Virginie Le Rolle. Model-based analysis of myocardial strains in left bundle branch block. Frontiers in Applied Mathematics and Statistics, 8, 2022.

[61] Bernard G. Krohn and Oscar Magidson. Left ventricular ejection time following cardioversion. The American Journal of Cardiology, 18(5):729–737, 1966.

[62] Morton E. Tavel, David O. Baugh, Harvey Feigenbaum, William K Nasser, and Janie Stewart. Left ventricular ejection time in atrial fibrillation. Circulation, 46(4):744–752, 1972.

[63] A. T. M. Gosselink, A. J. Smit, H. J. G. M. Crijns, H. H. Hillege, and K. I. Lie. Alteration of peripheral vasodilatory reserve capacity after cardioversion of atrial fibrillation. European Heart Journal, 17(6):926–934, 1996.

[64] Anton E Tuinenburg, Dirk J Van Veldhuisen, Frans Boomsma, Maarten P Van Den Berg, Pieter J De Kam, and Harry J.G.M Crijns. Comparison of Plasma Neurohormones in Congestive Heart Failure Patients With Atrial Fibrillation Versus Patients With Sinus Rhythm. The American Journal of Cardiology, 81(10):1207–1210, 1998.

[65] R. W. Stafford, W. S. Harris, and A. M. Weissler. Left Ventricular Systolic Time Intervals as Indices of Postural Circulatory Stress in Man. Circulation, 41(3):485–492, 1970.

[66] W Wieling, JJ Van Lieshout, and AD J Ten Harkel. Dynamics of Circulatory Adjustments to Head-Up Tilt and Tilt-Back in Healthy and Sympathetically Denervated Subjects. Clinical Science, 94(4):347–352, 1998.

[67] David Svec, Barbora Czippelova, Jana Cernanonva Krohova, Nikoleta Mazgutova, Radovan Wiszt, Zuzana Turianikova, Lenka Matuskova, and Michal Javorka. Short-Term Arterial Compliance Changes in the Context of Systolic Blood Pressure Influence. Physiological Research, 70(S3):S339–S348, 2021.

[68] Richard J Cohen, Ronald D Berger, and Theodore E Dushane. A quantitative model for the ventricular response during atrial fibrillation. IEEE Trans Biomed Eng, 30(12):769–781, 1983.

[69] Jie Lian, Dirk Müssig, and Volker Lang. Computer modeling of ventricular rhythm during atrial fibrillation and ventricular pacing. IEEE Trans Biomed Eng, 53(8):1512–1520, 2006.

[70] Valentina D.A. Corino, Frida Sandberg, Luca T. Mainardi, and Leif Sörnmo. An atrioventricular node model for analysis of the ventricular response during atrial fibrillation. IEEE Trans Biomed Eng, 58(12):3386–3395, 2011.

[71] M Masè, M Marini, M Disertori, and F Ravelli. Dynamics of AV coupling during human atrial fibrillation: role of atrial rate. Am J Physiol Heart Circ Physiol, 309(1):H198–H205, 2015.

[72] Tijmen Koopsen, Nick van Osta, Tim van Loon, Roel Meiburg, Wouter Huberts, Ahmed S. Beela, Feddo P. Kirkels, Bas R. van Klarenbosch, Arco J. Teske, Maarten J. Cramer, et al. Parameter subset reduction for imaging-based digital twin generation of patients with left ventricular mechanical discoordination. BioMedical Engineering OnLine, 23(46), 2024.

[73] Monica Maselli, Valter Giantin, Alessandro Franchin, Francesca Attanasio, Alessandra Tramontano, Pietro De Toni, Valentina Pengo, Domenico Corrado, and Enzo Manzato. Effect of restoring sinus rhythm in hypertensive patients with atrial fibrillation undergoing electrical cardioversion. Blood Pressure Monitoring, 21(6):335–339, 2016.

[74] Joakim Olbers, Ellen Jacobson, Fredrik Viberg, Nils Witt, Petter Ljungman, Mårten Rosenqvist, and Jan Östergren. Systolic blood pressure increases in patients with atrial fibrillation regaining sinus rhythm after electrical cardioversion. The Journal of Clinical Hypertension, 21(3):363–368, 2019.

[75] Joakim Olbers, Jan Östergren, Mårten Rosenqvist, Helga Skuladottir, Sofia Klavebäck, Petter Ljungman, and Nils Witt. Changes in 24-h ambulatory blood pressure following restoration of sinus rhythm in patients with atrial fibrillation. Journal of Hypertension, 39(2):243–249, 2021.

[76] J. R. Kieny, A. Sacrez, A. Facello, R. Arbogast, P. Bareiss, G. Roul, J.-L. Demangeat, B. Brunot, and A. Constantinesco. Increase in radionuclide left ventricular ejection fraction after eardioversion of chronic atrial fibrillation in idiopathic dilated cardiomyopathy. European Heart Journal, 13(9):1290–1295, 1992.

[77] Isabelle C. Van Gelder, Harry J.G.M. Crijns, Paul K. Blanksma, Martin L.J. Landsman, Jan L. Posma, Maarten P. Van Den Berg, Frits L. Meijler, and Kong I. Lie. Time course of hemodynamic changes and improvement of exercise tolerance after cardioversion of chronic atrial fibrillation unassociated with cardiac valve disease. The American Journal of Cardiology, 72(7):560–566, 1993.

[78] Hossein Sharifi, Lik Chuan Lee, Kenneth S. Campbell, and Jonathan F. Wenk. A multiscale finite element model of left ventricular mechanics incorporating baroreflex regulation. Computers in Biology and Medicine, 168:107690, 2024.

[79] Carlos Sánchez, Alfonso Bueno-Orovio, Esther Pueyo, and Blanca Rodríguez. Atrial Fibrillation Dynamics and Ionic Block Effects in Six Heterogeneous Human 3D Virtual Atria with Distinct Repolarization Dynamics. Frontiers in Bioengineering and Biotechnology, 5:29, 2017.

[80] Marta Varela, Ross Morgan, Adeline Theron, Desmond Dillon-Murphy, Henry Chubb, John Whitaker, Markus Henningsson, Paul Aljabar, Tobias Schaeffter, Christoph Kolbitsch, and Oleg V. Aslanidi. Novel MRI Technique Enables Non-Invasive Measurement of Atrial Wall Thickness. IEEE Transactions on Medical Imaging, 36(8):1607–1614, 2017.

[81] H Matsukubo, T Matsuura, N Endo, J Asayama, and T Watanabe. Echocardiographic measurement of right ventricular wall thickness. A new application of subxiphoid echocardiography. Circulation, 56(2):278–284, 1977.

[82] Jeroen Walpot, Daniel Juneau, Samia Massalha, Girish Dwivedi, Frank J. Rybicki, Benjamin J.W. Chow, and João R. Inácio. Left Ventricular Mid-Diastolic Wall Thickness: Normal Values for Coronary CT Angiography. Radiology: Cardiothoracic Imaging, 1(5):e190034, 2019.

[83] Daniel Burkhoff and John V Tyberg. Why does pulmonary venous pressure rise after onset of LV dysfunction: a theoretical analysis. American Journal of Physiology-Heart and Circulatory Physiology, 265:H1819–H1828, 1993.

[84] Attilio Reale. Acute Effects of Countershock Conversion of Atrial Fibrillation upon Right and Left Heart Hemodynamics. Circulation, 32(2):214–222, 1965.

